# Phylodynamic Analysis of the Global Dispersal and Spatiotemporal Dynamics of HIV-1 Subtype C

**DOI:** 10.64898/2026.06.03.729861

**Authors:** Xingguang Li, Sana Tamim, Gytis Dudas, Amanda Perofsky, Nídia S. Trovão

## Abstract

HIV-1 subtype C is the most prevalent HIV clade globally, yet its cross-continental transmission dynamics remain incompletely resolved due to sampling biases and unquantified analytical uncertainty. We performed a large-scale phylodynamic analysis of 1,221 near full-length genomic sequences from 32 countries (1986–2019). Employing a stratified subsampling strategy to mitigate bias, we systematically benchmarked 16 method combinations, pairing four maximum likelihood phylogenetic tools (FastTree, IQ-TREE, PhyML, RAxML-NG) with four temporal dating methods (TempEst, LSD, treedater, TreeTime). Topological analysis revealed significant discordance between rapid heuristics and higher-precision estimators. The estimated time to the most recent common ancestor (TMRCA) and the timing of individual introductions were highly sensitive to the temporal dating method, while evolutionary rates varied across both phylogenetic and temporal tools. Crucially, counts of viral introductions and the reconstruction of spatial transmission networks were statistically robust across all 16 combinations. Phylogeographic reconstruction confirmed a dominant, recurring dispersal corridor from Africa to Europe, with independent secondary reservoirs in South America and Asia. Ancestral trait reconstruction demonstrated transmission is primarily driven by heterosexual networks, within which “Not Recorded” risk groups were phylogenetically nested. This multi-method benchmarking reveals that while exact chronologies are sensitive to methodological choices, spatial network reconstructions and source-sink classifications remain remarkably consistent. These findings provide a robust map of subtype C’s global expansion and establish highly reliable, scalable workflows for resolving spatial dissemination routes for real-time genomic epidemiology.

## Introduction

Human immunodeficiency virus (HIV) remains one of the most profound global health challenges of the modern era. Since its emergence, the HIV pandemic has claimed 44.1 million lives, and by the end of 2024, approximately 40.8 million individuals were living with the virus worldwide, 65% of whom were in the World Health Organization (WHO) African Region^1^. While the rapid scale-up of antiretroviral therapy (ART) represents a monumental public health achievement, expanding from 7.7 million recipients in 2010 to 31.6 million by the end of December 2024^2^, the spread of HIV remains far from controlled. The financial implications of sustaining this response are staggering; modeling studies suggest that long-term financing needs for HIV control in sub-Saharan Africa alone could reach $261 billion by 2050 under scaled-up coverage scenarios^3^. This economic burden is further compounded by the virus’s vast genetic diversity^4^, which continues to challenge vaccine development, diagnostic sensitivity, and our understanding of transmission networks.

Among the four groups of HIV-1 (M, N, O and P), group M is responsible for the global pandemic and is further subdivided into subtypes (A–L)^5^. Of these, subtype C is the most prevalent lineage, accounting for the largest proportion of global infections^6^. Unlike subtype B, which predominates in the Americas and Western Europe, subtype C drives the severe generalized epidemics in Southern Africa, including South Africa, Botswana, and Zambia, as well as the large-scale epidemics in Ethiopia and India^7^. The evolutionary success of subtype C has been attributed to potential biological advantages, including higher replication capacity and viral loads across diverse host environments^8^, yet the precise historical pathways by which this clade spread from its origin to seed concurrent epidemics across three continents remain a subject of intense debate.

Reconstructing the history of such a widespread pathogen is facilitated by phylodynamic modeling. Phylogenetic analyses have established that the common ancestor of group M emerged in Kinshasa, Democratic Republic of the Congo (DRC), in the beginning of the 20^th^ century, with subtype C likely diverging in the mid-20^th^ century^9,10^. However, elucidating the subsequent dispersal events that drove its expansion out of Africa has proven challenging.

This challenge arises from two limitations in the existing literature. First, historical sampling bias has skewed our understanding of the pandemic. Genomic surveillance is often opportunistic, disproportionately favoring resource-rich countries in the Global North, while leaving substantial gaps in the regions where subtype C is most prevalent. As we demonstrated in our previous work, failing to account for these spatial and temporal disparities can lead to misleading conclusions about viral origins and migration rates^11^. Second, the field has largely relied on partial genomes^12^. Most HIV molecular epidemiology studies rely on short gene fragments (e.g., *pol* or *env*) generated for drug resistance testing. While cost-effective, these fragments often lack sufficient phylogenetic signal to resolve deep evolutionary nodes or distinguish rapid dissemination events. Furthermore, the choice of phylogenetic inference tool introduces an additional, unquantified layer of uncertainty. While Bayesian phylodynamic frameworks (such as BEAST^13–15^) offer a rigorous probabilistic approach to capturing this uncertainty, their high computational demands often render them impractical for real-time monitoring of large-scale genomic datasets, particularly in low- and middle-income settings^16^. Consequently, given the continuous growth of genomic surveillance data, analyses often rely on rapid, approximate likelihood heuristics (such as FastTree^17,18^), potentially sacrificing topological accuracy for computational speed. The extent to which these methodological shortcuts distort the reconstruction of global viral spread remains unknown.

To address these challenges, our research group has systematically developed a pipeline for high-resolution genomic epidemiology analyses. In our initial study^11^, we introduced a multiple-trait subsampling strategy designed to mitigate the inherent sampling biases in public databases. By subsampling sequences by location, time, and risk group simultaneously, we demonstrated that one could recover a more representative approximation of the true viral population structure. We subsequently applied this framework to characterize the emergence and circulation of subtype C, identifying key regional transmission hubs^19^.

In this study, we present our most comprehensive phylodynamic analysis of HIV-1 subtype C. Advancing from gene-fragment studies, we curated a dataset of 1,221 near full-length genomic sequences sampled from 32 countries between 1986 and 2019. To explicitly quantify and mitigate methodological uncertainty, we designed a combinatorial benchmarking experiment. We generated stratified subsampled datasets (ranging from 260 to 574 sequences) and analyzed them using 16 distinct phylogenetic-temporal method combinations, pairing four phylogenetic inference tools (FastTree^18^, IQ-TREE^20^, PhyML^21^, and RAxML-NG^22^) with four temporal dating methods (TempEst^23^, LSD^24^, treedater^25^, and TreeTime^26^).

By implementing this comprehensive benchmarking strategy, we quantify topological discordance between rapid heuristic methods and higher-precision estimators, using clustering information distance metrics and tanglegram visualizations to evaluate structural concordance. Concurrently, we evaluate the temporal sensitivity of these reconstructions by quantifying variance in estimates of the time to the most recent common ancestor (TMRCA) across alternative dating frameworks, thereby establishing a rigorous uncertainty interval for the subtype’s origin. Through this cross-validation, we seek to resolve the specific transmission networks driving intercontinental expansion and determine the stability of major transcontinental dispersal axes under varying model assumptions. We further assess the consistency of inferred transmission hubs and source-sink classifications across all method combinations, providing a quantitative framework for evaluating the robustness of phylogeographic conclusions.

## Materials and Methods

### Compilation and curation of HIV⍰1 subtype C genomes

All available near-complete HIV-1 subtype C genome sequences (corresponding to HXB2 positions 790–9417, with a minimum fragment length of 6,000 nt), with known sampling dates and geographic information, were retrieved from the Los Alamos National Laboratory (LANL) HIV Sequence Database^27^. Problematic sequences as defined by LANL were removed (including sequences with excessive non-ACTG characters indicative of laboratory contamination, artifactual deletions exceeding 100 nt, sequences shorter than 50 bp, or reverse-complement entries)^28^, and only one sequence per individual was retained prior to download. Sequence quality was assessed using the Quality Control tool^29^, genotype assignments were confirmed with RIP v3.0^30^, and hypermutation was evaluated using Hypermut v2.0^31^. The final dataset comprised 1,221 publicly available near-complete HIV-1 subtype C genome sequences (full1221), sampled between 1986 and 2019 and spanning 32 countries^11,19^. Sample origins were assigned using two-letter country codes^32^, and risk group metadata for sequences in full1221 were classified into six categories (risk6): men who have sex with men (MSM), persons who inject drugs (PI), heterosexual (SH), mother–baby (MB), not recorded or unknown (NR), and remaining sequences were assigned to ‘other’ (OT).

### Stratified subsampling for representative datasets

Multiple sequence alignments of the full1221 dataset were generated using MAFFT v7.427^33^ under default settings and subsequently adjusted manually in BioEdit v7.2.5^34^. Subsequently, sequences with >50% gaps, lengths <99% of the multiple sequence alignment, or duplicates, defined by identical sampling date, country, risk group, and nucleotide sequence, were excluded. This resulted in a full-genome dataset comprising 1,098 sequences (full1098).

Our previous studies showed that incorporating as many traits as possible during subsampling produces more balanced datasets that capture the overall circulating viral diversity while reducing potential biases in the original data^11,19^. Therefore, further subsampling was performed using SAMPI (https://github.com/jlcherry/SAMPI)^35^ to generate balanced datasets comprising 2, 4, 6, 8, or 10 sequences per country, risk group, and sampling date, with each subsampling replicated three times. This resulted in full-genome datasets comprising 260, 378, 468, 527, and 574 sequences, subsampled by sampling location, risk group, and collection date (locrisk260.1–3, locrisk378.1–3, locrisk468.1–3, locrisk527.1–3, and locrisk574.1–3).

### Model selection and phylogenetic inference

The best-fit nucleotide substitution model was identified from 24 candidate models using ModelTest-NG v0.2.0^36^. The general time-reversible (GTR) substitution model with a FreeRate component (GTR+R4) was determined to be the best-fit model for all datasets according to the Akaike Information Criterion (AIC), corrected AIC (AICc), and Bayesian Information Criterion (BIC). To evaluate the phylogenetic signal in each dataset, likelihood-mapping analysis was performed using IQ-TREE v2.3.6, based on 10,000 randomly selected quartets under the GTR+R4 model^37^. Pythia v2.0.0 was used to assess the difficulty of phylogenetic inference for each dataset prior to setting up and conducting phylogenetic tree reconstruction^38^. Maximum-likelihood (ML) phylogenies were inferred using FastTree v2.1.11^18^, IQ-TREE v2.3.6^20^, PhyML v3.3.20220408^21^, and RAxML-NG v1.2.2^22^ under the GTR+Γ4 model (for FastTree, which does not support FreeRate models) or the GTR+R4 model (for IQ-TREE, PhyML, and RAxML-NG).

### Topological comparison and visualization

To rigorously evaluate the consistency of phylogenetic reconstructions across the four inference tools, we applied both quantitative and qualitative approaches. First, rather than using the standard Robinson-Foulds metric^39^, which is highly sensitive to minor leaf instability, we calculated the Clustering Information Distance (CID) between tree topologies using the TreeDist R package^40^. This information-theoretic metric quantifies the distance between trees in bits, reflecting the extent to which the splits of one tree inform those of another. Lower CID values indicate greater shared information and stronger topological concordance.

Second, to visually inspect these structural discrepancies, we generated tanglegrams using the baltic Python library^41^. Tanglegrams were generated by pairing the best-scoring ML trees inferred from identical subsampled datasets, such as replicates from FastTree and RAxML-NG. Trees were rotated to reduce entanglement, minimizing crossing lines between corresponding taxa. This allowed the identification of specific clades or lineages that were inconsistently resolved by approximate heuristics relative to higher-precision algorithms.

### Evolutionary dynamics

Temporal signal analysis and outlier detection were evaluated using TempEst v1.5.3^23^. Evolutionary rates (ER) and (TMRCAs) for ML phylogenies were estimated using four approaches: root-to-tip regression in TempEst v1.5.3^23^, least-squares dating in LSD v.2.4.1^24^, Gamma-Poisson mixture model in treedater^25^ and maximum-likelihood dating in TreeTime v0.11.4^26^. Each method yielded estimates of the ER (substitutions per site per year) and TMRCA, with uncertainty assessed from variation across methods. Comparison across these independent approaches enabled assessment of the robustness of the estimates.

### Phylogeographic and risk group reconstruction

To infer the spatial and demographic history of the virus, we employed discrete-trait ancestral (DTA) reconstruction using the mugration model implemented in TreeTime v0.11.4^26^. This analysis was performed on all time-scaled topologies generated by the four dating methods. To assess the sensitivity of our findings to risk group classification, we employed a hierarchical grouping strategy. We first analyzed the full six-category scheme (Risk6: SM, PI, SH, MB, NR, OT). Subsequently, we performed sensitivity analyses by merging under-sampled groups: first merging NR into OT to create a 5-level scheme (Risk5), then merging MB into OT to generate a 4-level scheme. Ancestral states were reconstructed for both geographic location (continent, region, country) and risk group at all internal nodes.

### Statistical Benchmarking of Phylodynamic Methods

To assess the impact of methodological choices on phylodynamic inference, we performed a statistical validation across 16 method combinations (4 phylogenetic x 4 temporal methods) using the 5 subsampled datasets with 3 replicates each. We evaluated the variance in three key metrics: TMRCA, ER, and the number of viral introduction events.

We used Kruskal-Wallis rank sum tests to assess differences in TMRCA, ER, and introduction count estimates across temporal methods, phylogenetic reconstruction methods, and subsampled datasets. Effect sizes for Kruskal-Wallis tests were quantified as the eta-squared statistic (’}²), with values interpreted as small (’}² < 0.06), moderate (’}² = 0.06 – < 0.14), or large (’}² > 0.14). When Kruskal-Wallis tests were statistically significant (p < 0.05), we performed pairwise comparisons using post-hoc Dunn tests with Bonferroni correction. To explicitly test for interdependence between the choice of method for phylogenetic reconstruction and temporal dating, we employed the Scheirer-Ray-Hare test^42^, a non-parametric two-way factorial analysis of variance on rank-transformed data. We measured variability in TMRCA, ER, and introduction counts across subsampling replicates and method combinations using the coefficient of variation (CV).

To further assess the sensitivity of inferred introduction times to methodological choices, we fit linear mixed-effects models (LMMs) to individual introduction time estimates. The response variable was the estimated introduction time (decimal year) for each inter-country transmission event. Fixed effects included the temporal dating method, the phylogenetic reconstruction method, and their interaction. To account for the hierarchical structure of the data, random intercepts were specified for origin country, destination country, transmission route (origin-destination country pair), and subsampling replicate. To ensure balanced comparisons, we retained only country pairs detected by all 16 method combinations within each dataset (pairs did not need to appear in every dataset, as smaller subsamples naturally detect fewer routes). We assessed the significance of fixed effects using Type III analysis of variance (ANOVA) with Satterthwaite’s degrees of freedom approximation. We tested for differences in the temporal distribution of introduction events across methods using chi-squared tests of independence. Model robustness was evaluated through alternative random effects structures, rank-transformed LMM sensitivity analyses, and non-parametric Kruskal-Wallis tests.

To determine whether the identification of transmission hubs is robust to methodological choices, we used Kendall’s coefficient of concordance (W)^43^ to evaluate the concordance of StrainHub degree centrality rankings across phylogenetic methods and subsampled datasets. For each geographic resolution (continent, region, country) and risk group classification scheme (risk4, risk5, risk6), W was calculated for each centrality metric to test whether the ranking of geographic locations or risk groups was consistent across raters. W ranges from 0 (no agreement) to 1 (perfect agreement), with values above 0.7 indicating strong concordance. Three rater groupings were evaluated: phylogenetic reconstruction method (k = 4, averaged across datasets and subsampling replicates), dataset size (k = 5, averaged across ML methods and replicates), and all method-dataset combinations (k = 60, treating each ML method × dataset × replicate as an independent rater). Statistical significance was assessed using the Friedman chi-squared approximation.

To assess whether source-sink classifications derived from DTA reconstruction were similarly robust to methodological choices, we applied the same concordance framework to the Source-Sink Score (SSS) and its components, export and import counts, across the same resolution levels (continent, region, country, risk4, risk5, risk6)^44^. As described above, DTA reconstruction was performed on time-scaled trees from all four temporal dating methods, yielding source-sink scores for all 16 method combinations. The rater space was therefore expanded relative to the StrainHub analysis to include temporal dating method as an additional dimension, yielding four rater groupings: phylogenetic method (k = 4), temporal dating method (k = 4), dataset size (k = 5), and all method-dataset combinations (k = 240, treating each temporal method × ML method × dataset × replicate as an independent rater). To ensure concordance estimates reflected observed rankings only, locations were retained for a given dataset only if detected by all 16 method combinations within that dataset; locations not meeting this criterion were excluded rather than imputed. Locations did not need to appear in every dataset, as smaller subsamples naturally detect fewer nodes. At the country level, two locations (Cyprus, United Kingdom) were excluded from the analysis, as both were detected in only one method combination within the locrisk574 dataset and were absent from all smaller subsamples.

Benchmarking analyses were conducted in R (version 4.5.2)^45^ using the rstatix^46^, rcompanion^47^, lme4^48^ and lmerTest^49^ packages.

## Results

### Global distribution and dataset composition

To characterize the global diversity of HIV-1 Subtype C while mitigating the severe sampling biases inherent in public repositories^27^, we analyzed near full-length genomic sequences sampled from 32 countries between 1986 and 2019^11,19^. From the initial pool of available genomes, we generated five stratified datasets of increasing size (260, 378, 468, 527, and 574 sequences per dataset). Each dataset was generated in triplicate (replicates 1–3) to ensure that downstream phylodynamic inferences were not artifacts of random sampling stochasticity (Figure 1). This resulting collection of 15 analyzed datasets covers six major risk groups and provides a balanced spatiotemporal representation of the pandemic, forming the basis for the subsequent methodological benchmarking (Supplementary Figures S1–S2).

**Figure 1.**
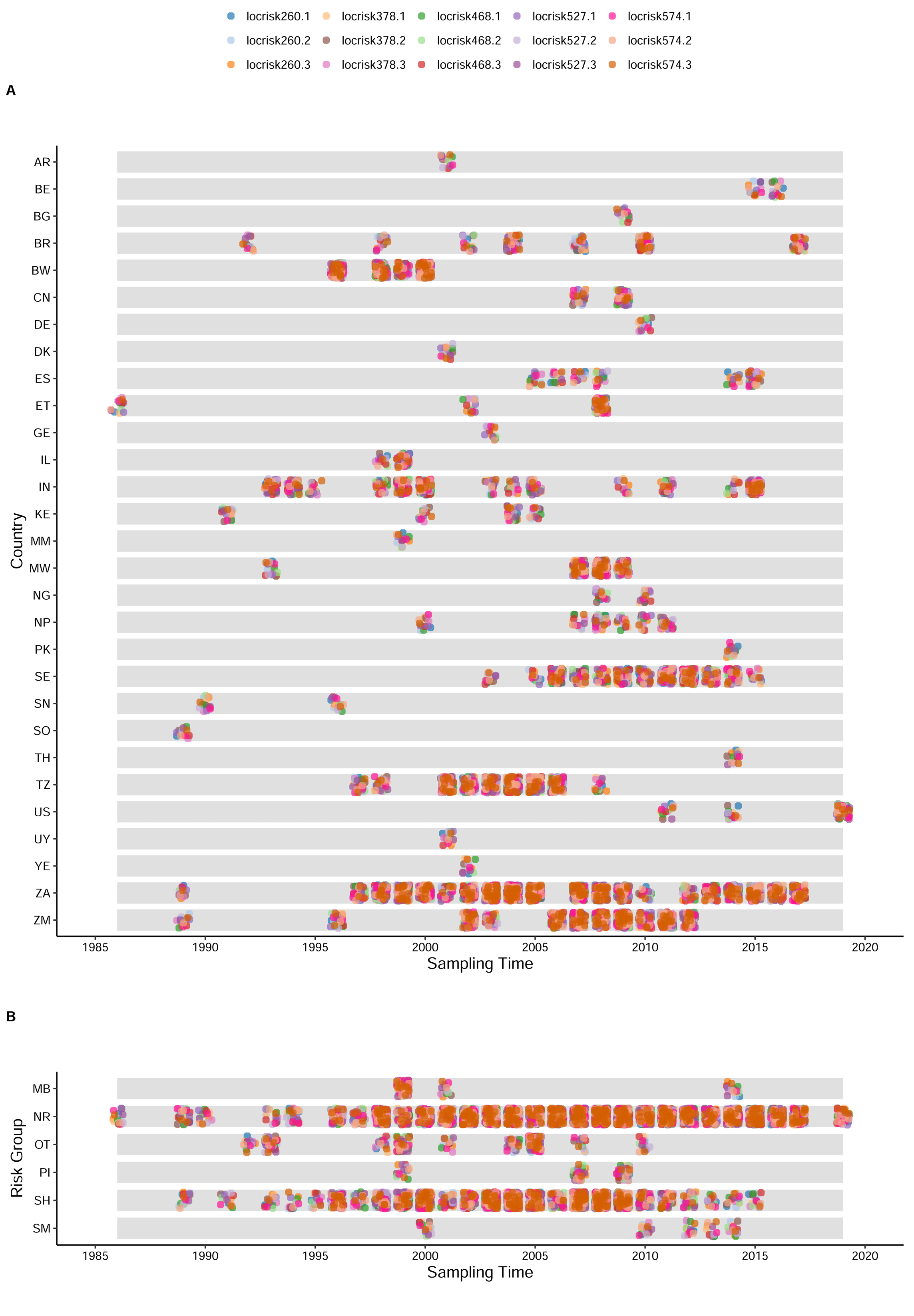
Temporal distribution of fifteen sub-sampled datasets of HIV subtype C genomes used in this study. (A) Stratified by country. (B) Stratified by risk group.

### Phylogenetic Signal and Temporal Resolution

Before performing complex phylodynamic reconstruction, we assessed the information content and temporal structure of the subsampled datasets. Likelihood-mapping analysis^37^ revealed a strong phylogenetic signal across all replicates; most sequence quartets (92.2–93.1%) were distributed at the corners of the likelihood triangle, rather than at the center (5–6%) or along the sides (6.3–7.2%), indicating a fully resolved, tree-like evolutionary history with minimal phylogenetic noise (Supplementary Figure S3).

Temporal signal analysis using TempEst confirmed the presence of a molecular clock (Figure 2). Linear regression of root-to-tip genetic distances against sampling dates demonstrated a strong temporal signal across all ML topologies (FastTree, IQ-TREE, PhyML, and RAxML-NG), with correlation coefficients ranging from 0.55 to 0.58 (p < 0.0001). Visual inspection of the regression plots revealed no significant outlier sequences that would distort evolutionary rate estimates. This robust temporal structure validates the suitability of these datasets for molecular clock dating, providing a solid foundation for the comparative TMRCA and dispersal analyses detailed below. We additionally applied Pythia^38^ to the subsampled alignments. Difficulty scores for phylogenetic inference increased with subsample size, ranging from 0.34–0.36 in the locrisk260 datasets to 0.58–0.59 in the locrisk574 datasets, with minimal variation among replicate datasets within each size class. All subsampled alignments fell within the intermediate difficulty category (Supplementary Table S1).

**Figure 2.**
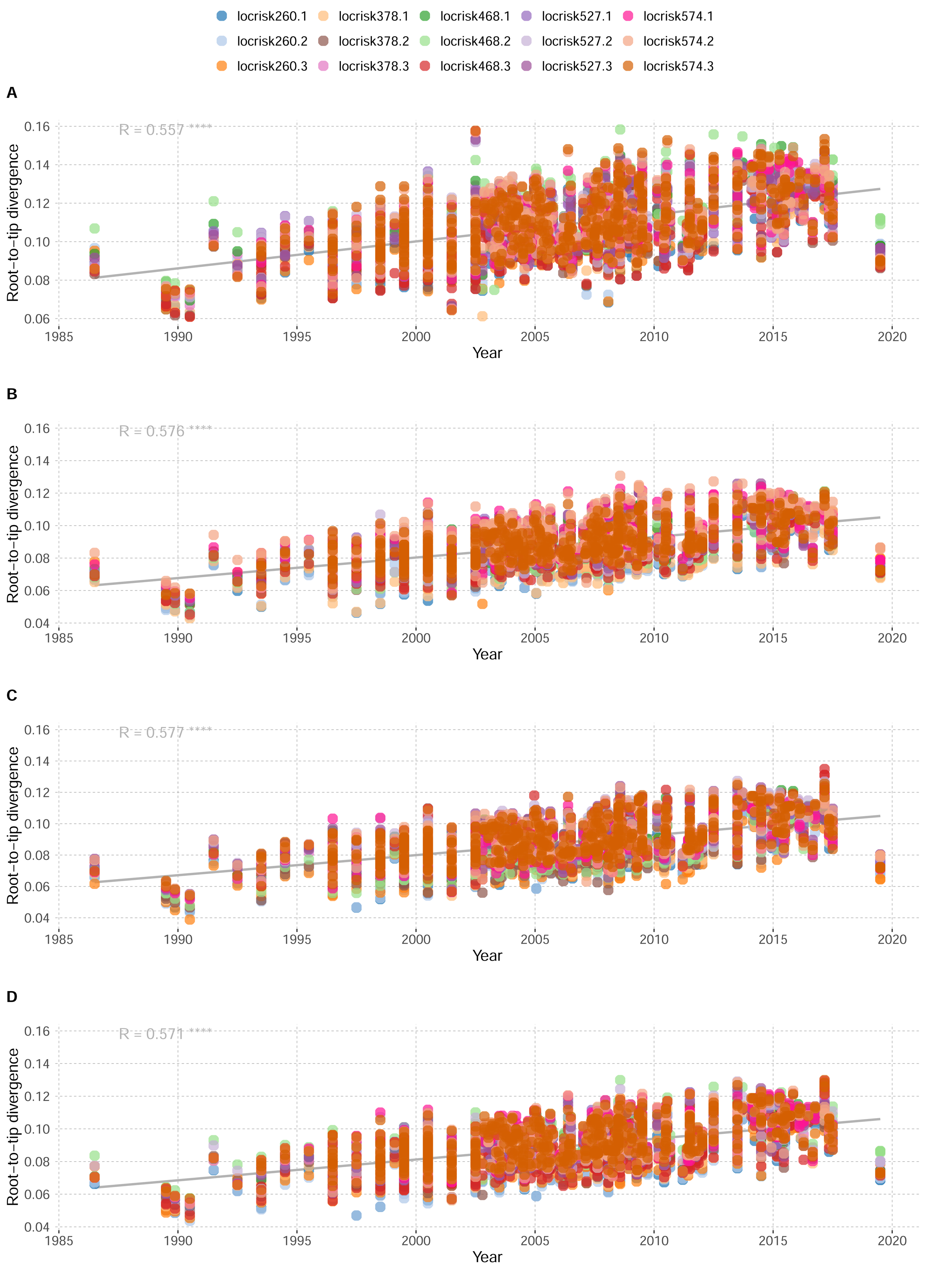
Root-to-tip regression analyses illustrating the relationship between genetic divergence and sampling time across fifteen sub-sampled HIV-1 subtype C datasets, grouped by phylogenetic method. (A) FastTree, (B) IQ-TREE, (C) PhyML, and (D) RAxML-NG.

### Topological Congruence and Methodological Validation

To ensure the robustness of our phylodynamic inferences, we systematically evaluated topological consistency across four ML inference tools. Quantitative analysis of tree space using Clustering Information Distance (CID) revealed significant topological discordance between rapid heuristic methods and higher precision algorithms (Figure 3). Specifically, trees inferred by FastTree exhibited the highest divergence from other methods, with a mean information distance of 37.9 (vs. PhyML) and 36.5 (vs. RAxML-NG). In contrast, the higher precision estimators converged on more stable topologies; the highest congruence was observed between PhyML and RAxML-NG (mean distance: 23.2), followed closely by IQ-TREE comparisons. Visual inspection of tanglegrams (Figure 4 and Supplementary Figures S4-S17) corroborated this quantitative trend, identifying FastTree as a consistent topological outlier across all subsampling replicates. To assess whether these topological differences propagated into downstream phylodynamic estimates, we performed statistical validation across all method combinations. While phylogenetic method had no significant impact on TMRCA estimates (Kruskal-Wallis χ²(3) = 5.59, p = 0.13, η² = 0.011), it significantly affected ER estimates (χ²(3) = 21.9, p < 0.0001, η² = 0.08) (Supplementary Figure S18). Post-hoc comparisons revealed that FastTree produced significantly higher evolutionary rates than the three higher-precision methods (FastTree vs. IQ-TREE, PhyML, RAxML-NG: all adjusted p < 0.01), while IQ-TREE, PhyML, and RAxML-NG were statistically equivalent (all adjusted p > 0.05) (Supplementary Figure S18).

**Figure 3.**
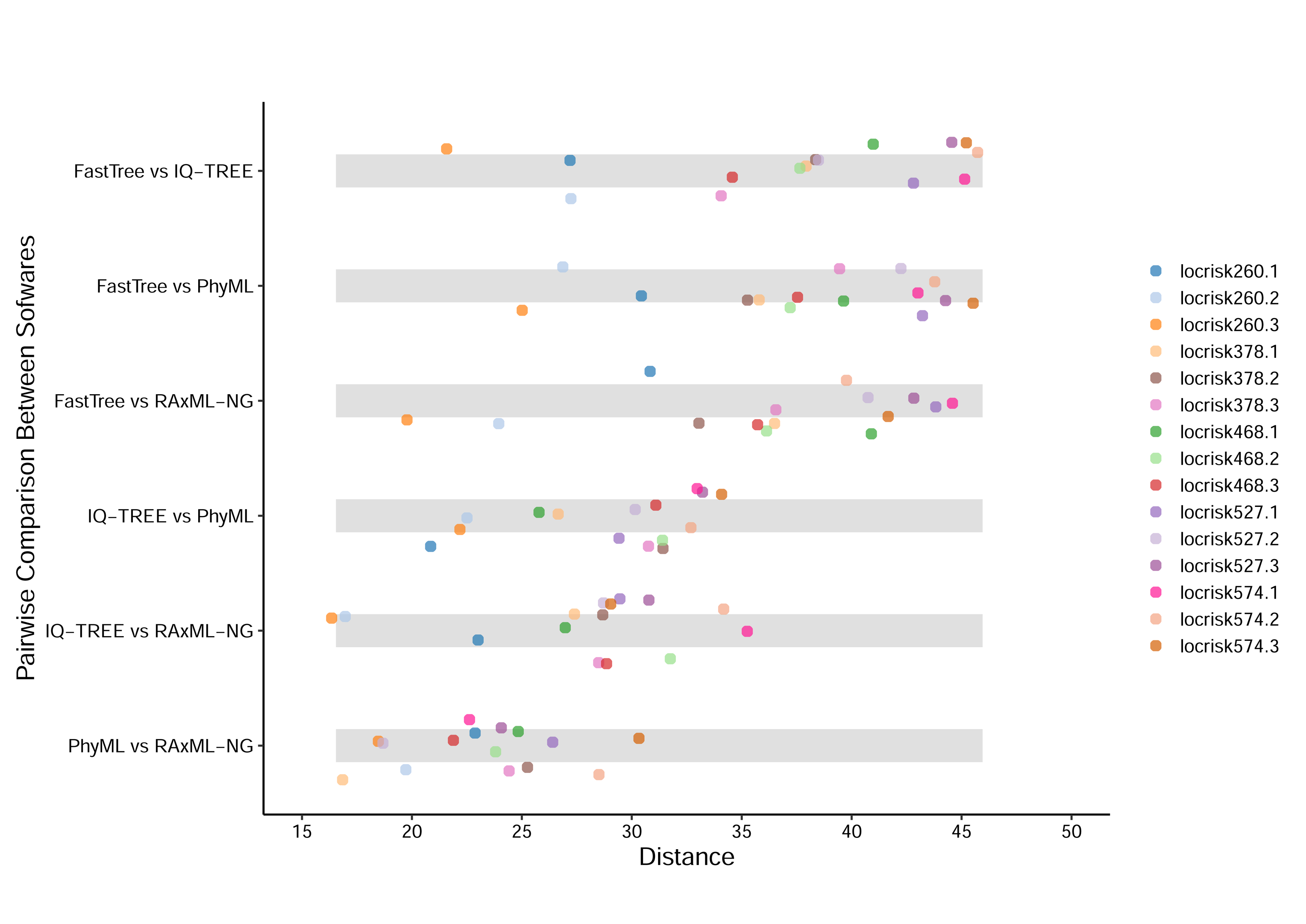
Pairwise comparisons of phylogenetic tree distances among four phylogenetic softwares (FastTree, IQ-TREE, PhyML, and RAxML-NG) across fifteen sub-sampled HIV-1 subtype C datasets, calculated using TreeDist.

**Figure 4.**
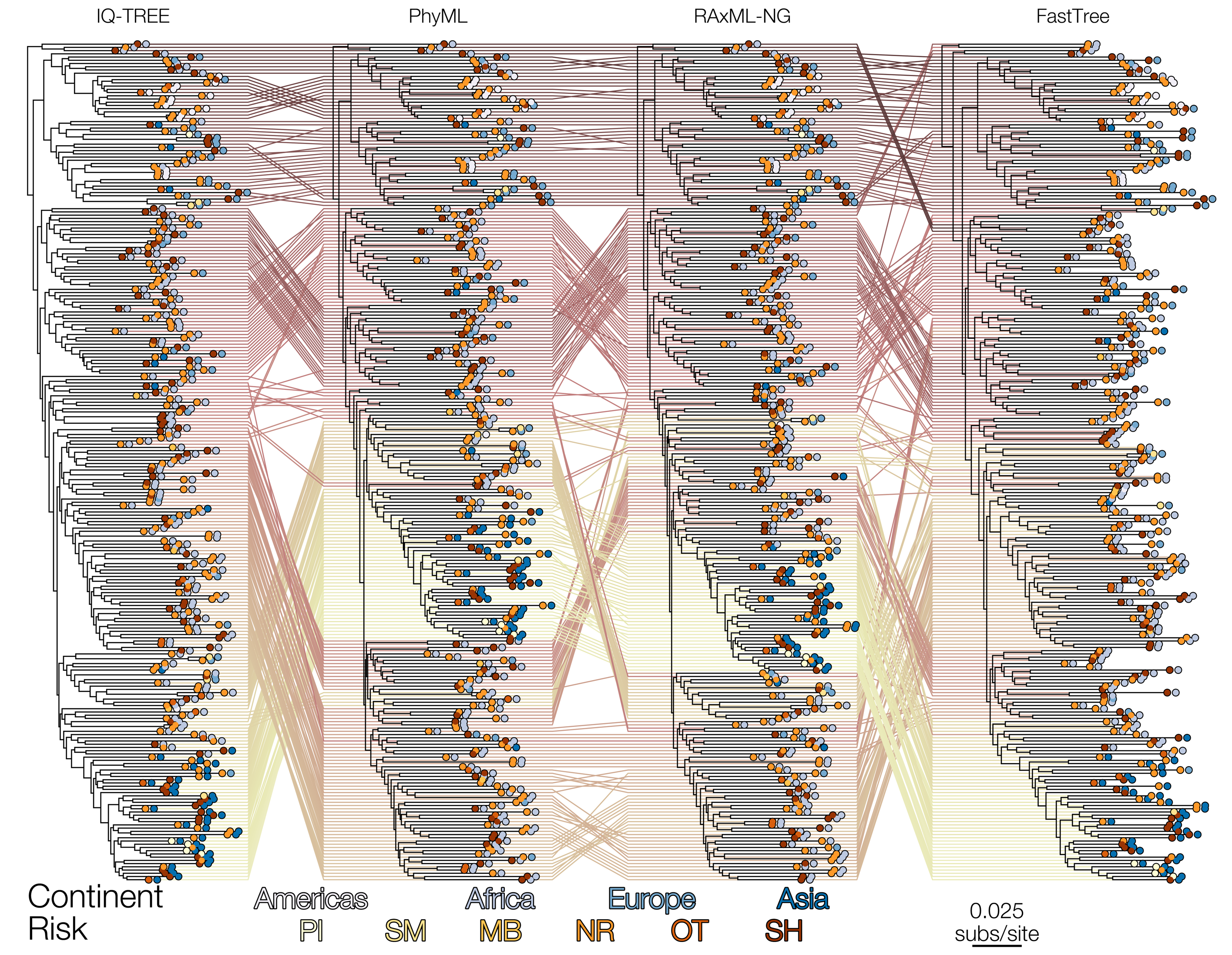
Tanglegram of phylogenetic trees inferred by four phylogenetic softwares ordered by average-linkage clustering based on ClusteringInfoDistance for locrisk260.1. Tangle lines are coloured according to the tip order along the y-axis of the leftmost tree. Each tip is annotated with a pair of circles representing risk group (left) and continent (right), as indicated in the legend. This is one of fifteen sub-sampled HIV-1 subtype C datasets, which can be found in Supplementary Figures S4-S17.

### Robustness of Phylodynamic Estimates

Statistical benchmarking of 16 method combinations revealed that the choice of inference tool significantly impacts temporal estimates but not spatial reconstruction. Estimates of the TMRCA and ER were significantly influenced by the choice of temporal dating method (Kruskal-Wallis χ²(3) = 119.9, p < 0.0001, η² = 0.495 for TMRCA; χ²(3) = 143.7, p < 0.0001, η² = 0.596 for ER) (Supplementary Figure S18). All pairwise comparisons between temporal methods were statistically significant for TMRCA (post-hoc Dunn’s tests, all adjusted p < 0.05), resulting in an 18-year interval of uncertainty between the earliest (TreeTime: mean 1938) and latest (treedater: mean 1955) estimates. For evolutionary rates, TempEst and treedater produced statistically equivalent estimates (post-hoc Dunn’s tests, adjusted p = 0.63), while all other temporal method pairs differed significantly (all adjusted p < 0.05).

As noted above, the method for phylogenetic reconstruction significantly impacted ER estimates (Kruskal-Wallis χ²(3) = 21.9, p < 0.0001, η² = 0.08) but not TMRCA (χ²(3) = 5.59, p = 0.13, η² = 0.011), with FastTree producing higher rates than the three higher-precision methods. This trend is clearly visible in the heatmaps of mean ER across method combinations (Supplementary Figure S19). We did not detect significant interactions between phylogenetic and temporal methods for TMRCA (Scheirer-Ray-Hare test, H(9) = 7.15, p = 0.62) or ER (H(9) = 14.02, p = 0.12).

Consistency across replicates varied by dataset size and method combination. The smallest subsampled dataset (260 sequences) produced the most consistent estimates for both TMRCA (coefficient of variation, mean CV = 0.1%) and ER (mean CV = 1.72%). Among method combinations, LSD paired with RAxML-NG yielded the most consistent TMRCA estimates (mean CV = 0.06%), while LSD paired with IQ-TREE produced the most consistent ER estimates (mean CV = 0.89%). Overall, CV was low across all method combinations (TMRCA: 0.06 – 0.25%; ER: 0.89 – 8.83%), indicating good reproducibility.

The reconstruction of viral dispersal patterns was highly robust to these temporal fluctuations. The mean number of inferred inter-continental introduction events did not differ significantly across temporal methods (Kruskal-Wallis χ(3)² = 1.31, p = 0.73, η² = - 0.001), phylogenetic methods (χ²(3) = 0.11, p = 0.99, η² = -0.002), or their interaction (Scheirer-Ray-Hare test, H(9) = 0.82, p > 0.99) (Supplementary Figure S18). TreeTime paired with IQ-TREE produced the most consistent introduction count estimates (mean CV = 11.8%), with all method combinations showing relatively low variation (CV range: 11.8–16.4%) (Supplementary Figures S18-S19). This indicates that the dispersal patterns of HIV-1 Subtype C represent a strong biological signal resilient to methodological variation.

### Spatiotemporal Dispersion

Based on the validated higher precision topologies (RAxML-NG/PhyML) and the preferred temporal signal resolution (treedater), we reconstructed the evolutionary history of HIV-1 Subtype C. To provide the highest resolution of transmission dynamics, we present findings from the largest stratified dataset (locrisk574); results from all other subsampling replicates yielded congruent patterns and are detailed in Supplementary Materials.

The time to the most recent common ancestor (TMRCA) for this dataset was estimated between 16 October 1938 and 6 November 1951 (Table 3), consistent with the consensus emergence of the subtype in the mid-20th century. The estimated evolutionary rate was approximately 1.16 x 10^-3^ to 1.49 x 10^-3^ substitutions per site per year.

Phylogeographic reconstruction identified Southern Africa as the primary hub of global dissemination. We identified a total of 46–52 distinct inter-continental transmission events. Of note, global dispersal was highly directional: the majority of inter-continental transitions occurred from Africa to Europe, driven by multiple independent introduction waves rather than a single founder event. Secondary dissemination routes were observed from Africa to South America (specifically Brazil) and Asia, though these represented a smaller fraction of the global transmission network.

### Robustness of Phylogeographic Inferences

Statistical benchmarking confirmed the robustness of these phylogeographic inferences across methodological choices. Centrality rankings derived from StrainHub showed strong to very strong concordance across all geographic resolution levels and all 60 method-dataset combinations (continent: mean Kendall’s W = 0.85, range 0.80–0.95; region: mean W = 0.80, range 0.70–0.86; country: mean W = 0.80, range 0.71–0.91; all p < 0.0001; Supplementary Table S2). Africa was consistently identified as the primary source hub and Europe as the principal recipient, and South Africa, Sweden, and Zambia consistently ranked as the most central countries in the global transmission network regardless of method or dataset.

Source-sink scores (SSS) derived from DTA reconstruction were similarly robust across geographic resolution levels and all 240 method-dataset combinations (all p < 0.0001; Supplementary Table S3). Concordance was very strong at the continent level (mean W = 0.93, range 0.90–0.95) and strong at the region and country levels (region: mean W = 0.84, range 0.75–0.95; country: mean W = 0.74, range 0.68–0.85), with import concordance consistently higher than export and SSS at the country level (W = 0.85 vs. 0.68 and 0.70), indicating that sink country identification is more stable across methods than source identification. Concordance was similarly high across phylogenetic and temporal dating methods evaluated separately (mean W = 0.88–0.98 for both), confirming that source-sink classifications are robust to both dimensions of methodological choice.

### Sensitivity of Introduction Timing to Methodological Choice

As noted above, the total number of introduction events was stable across temporal methods (Kruskal Wallis χ(3)² = 1.31, p = 0.73), phylogenetic methods (χ²(3) = 0.11, p = 0.99), and their interaction (p > 0.99), with consistently low variation across all method combinations (CV range: 11.8–16.4%). However, the inferred timing of individual introduction events was sensitive to the choice of temporal dating method. We fit a linear mixed-effects model to individual introduction time estimates across all shared dispersal routes (n = 24,781 events across 58 country pairs; Table S4), revealing a significant effect of temporal dating method (ANOVA, F = 108.55, p < 0.0001) and phylogenetic method (F = 3.05, p = 0.027), but no significant interaction between these two factors (F = 1.4, p = 0.18). Although the effect of phylogenetic method was statistically significant, we did not observe directional patterns in the timing of introductions (i.e., consistently earlier or later timing) across methods, suggesting differences in route-specific topology rather than systematic bias.

Among temporal methods, TempEst estimated introduction times 6.3 years earlier than the reference category, LSD (95% CI: −7.6 to −5.0, p < 0.001), while TreeTime estimated introductions 1.6 years earlier than LSD (95% CI: −2.9 to −0.3, p = 0.016); treedater did not differ significantly from LSD (0.1 years; 95% CI: −1.2 to 1.4, p = 0.87). This temporal disagreement was concentrated in early (pre-1980) divergence events, wherein TempEst estimated substantially earlier introduction dates than other methods. For introductions after 1980, temporal methods converged, agreeing within approximately 1 year for the 1980–2000 period and within 0.5 years after 2000. Notably, temporal dating methods also differed in the general timing of waves of introductions: when aggregating across all dispersal routes, LSD, TreeTime, and treedater reconstructed two distinct waves separated by a clear valley (∼1975–1985), whereas TempEst inferred a significantly earlier first wave and more continuous pattern of low-level dispersal during the intervening period between waves (χ² = 159.53, df = 6, p < 0.0001) (Supplementary Figure S20).

Variance decomposition indicated that the largest source of variation in introduction timing was the specific dispersal route rather than methodological choice, with destination country accounting for 22.6% of variance, origin country for 4.8%, and country pair for 9.2%, while subsampling replicate contributed negligibly (< 0.1%). Despite its statistical significance, methodological choice explained less than 1% of variance in introduction timing (marginal R² = 0.009), while the random effects structure, which captures systematic differences among dispersal routes and geographic origins/destinations, accounted for 37% (intraclass correlation coefficient (ICC) = 0.37). These conclusions were robust to alternative random effects structures, rank-transformed sensitivity analyses, and non-parametric tests.

### Risk Group Dynamics

Ancestral trait reconstruction of risk groups revealed that heterosexual contact (SH) was the primary driver of global mobility. The ‘SH’ risk group functioned as the major source population for transitions into other demographic categories. Notably, sequences classified as ‘Not Recorded’ (NR) consistently clustered within established heterosexual clades across all subsampling replicates.

When assessing concordance of StrainHub centrality rankings for risk groups across phylogenetic methods and datasets, out-degree rankings were highly concordant across all classification schemes (W = 0.81–0.89), confirming robust identification of source populations. However, in-degree concordance was notably lower across all 60 method-dataset combinations (W = 0.17–0.40), indicating that sink population identification is less stable across methods, an asymmetry not observed for geographic levels, where in-degree concordance was strong across all resolutions (W = 0.71–0.91; Supplementary Table S2). Source-sink scores derived from DTA reconstruction showed similarly strong concordance for risk group source identification across all three classification schemes and all 240 method-dataset combinations (risk6: mean W = 0.83; risk5: mean W = 0.82; risk4: mean W = 0.70; all p < 0.0001), with heterosexual contact consistently ranking as the dominant source group regardless of risk group classification scheme (Supplementary Table S3).

## Discussion

This study benchmarks 16 phylodynamic model combinations across 1,221 near full-length genomes to resolve the evolutionary history of HIV-1 Subtype C, distinguishing biological signal from methodological variance. Results indicate that while TMRCA and ER are sensitive to methodological choices, yielding an 18-year interval of uncertainty for the subtype’s origin, the reconstruction of spatial transmission networks remains statistically stable across methods. We identify a persistent transmission axis from Africa to Europe, driven primarily by heterosexual networks. These findings align with previous work on sampling bias^50^ and general emergence^51^, confirming that the Africa-to-Europe dispersal corridor is a consistent feature of the HIV pandemic’s history rather than an artifact of tree inference or sampling stochasticity.

Quantification of phylodynamic uncertainty reveals significant performance disparities among inference tools. Rapid heuristic methods (FastTree) produced topologies with high divergence from precision estimators (Clustering Information Distance > 36), supporting their exclusion from detailed transmission studies. Additionally, the estimated TMRCA of subtype C varied depending on the dating algorithm, ranging from the 1930s (TreeTime) to the 1950s (treedater). This variance implies that single-method maximum likelihood studies may underestimate confidence intervals for viral origins. Our findings regarding the sensitivity of time-dependent parameters to the choice of dating algorithm align with recent work by Fourment et al. (2025)^63^, which demonstrated that fixing tree topologies biases temporal estimates compared to fully Bayesian inference. However, our analysis extends this evaluation to demonstrate that the spatial and demographic reconstruction of transmission networks remains highly statistically stable (p > 0.73 across methods) despite this temporal variance. This indicates that while fixed-topology heuristics may introduce chronological uncertainty, they remain highly reliable, scalable tools for resolving global dissemination routes and source-sink dynamics in real-time genomic epidemiology. Concordance analyses of both network centrality rankings and source-sink scores further confirmed this stability, with strong to very strong agreement across geographic resolution levels regardless of phylogenetic or temporal method choice.

Extending this analysis to individual introduction events, we found that temporal dating methods also systematically shift the estimated timing of dispersal events. After accounting for geographic route and replicate effects, TempEst placed introductions approximately 6 years earlier than LSD (the reference category in regression analyses). Critically, the methods also differed in the inferred temporal structure of introductions: LSD, treedater, and TreeTime reconstructed two distinct introduction waves, while TempEst inferred a much earlier first wave and more continuous dispersal during the intervening period. This discrepancy has implications for the biological narrative of the HIV pandemic, as discrete seeding events versus sustained transmission reflect different underlying epidemiological dynamics. The disagreement was concentrated in early divergence events (pre-1980), while post-1980 introductions were dated consistently across all methods.

Phylogeographic analysis indicates highly directional global dissemination. The data show direct, repeated introductions from Southern Africa to Europe. This dispersal pattern likely reflects deep historical connectivity and economic- and tourism-associated human mobility^52–54^, facilitating the introduction of the subtype into European networks. Quantitatively, Europe represented the primary sink for inter-continental dispersal, accounting for the majority of all inferred export events outside of Africa. Within Europe, we identified frequent viral introductions into Sweden, Spain, and Belgium, likely reflecting both the intensity of the migration pressure and the density of available surveillance data in these nations^55,56^.

Beyond this primary axis, we observed significant secondary dissemination routes to Asia and South America. In Asia, India emerged as the dominant secondary reservoir^57^, receiving the highest number of introductions outside of Europe, followed by distinct lineages established in China and Nepal^58^. Additionally, Israel represented a unique and high-frequency destination, with introduction counts comparable to major European hubs^59^, suggesting a specific and active transmission corridor distinct from the broader Asian epidemic.

In South America, dissemination was characterized by a cluster with balanced introduction events identified in Brazil, Argentina, and Uruguay^60,61^. Unlike the high-frequency re-seeding observed in Europe, these South American lineages suggest a pattern of successful local establishment following fewer founder events. The detection of these concurrent, yet geographically distinct, dispersal waves highlights the complexity of subtype C’s global expansion, in which Europe acts as a major “sink” for sustained, high-volume introductions, while South America and Asia represent independent reservoirs where the virus has successfully established local sub-epidemics adapted to regional networks.

Ancestral trait reconstruction provides resolution for the “Not Recorded” (NR) risk category, which constitutes a significant proportion of public database entries and often confounds epidemiological modeling. Our analysis reveals that the phylogenetic placement of NR sequences is non-random. Unlike “Men who have Sex with Men” (MSM) and “People who Inject Drugs” (PWID), which typically form distinct, monophyletic transmission chains indicative of compartmentalized sub-epidemics^62^, NR sequences were diffusely distributed throughout the topology and consistently nested within major Heterosexual (SH) lineages. This pattern provides strong evolutionary evidence that the NR category is not a mixture of undisclosed high-risk behaviors (which would likely cluster with MSM or PWID clades) but is systematically dominated by heterosexual transmission. This misclassification implies that current surveillance metadata obscures the true extent of the heterosexual epidemic, as a substantial fraction of “unknown” transmission represents the background of the generalized heterosexual epidemic rather than cryptic key populations^51^.

Importantly, risk group source-sink assignments were robust to methodological choice: out-degree centrality rankings (reflecting source population identification) showed strong concordance across all method combinations and risk group classification schemes, while in-degree concordance was notably lower, suggesting that sink population identification is inherently less stable across methods. This asymmetry suggests that source populations are more reliably identified across methods than sink populations, and that conclusions about recipient risk groups should be interpreted with greater caution in single-method analyses.

This study has several limitations. First, despite stratified subsampling, the analysis is constrained by the availability of full-genome sequences, which are sparse for certain high-prevalence regions relative to pol fragments. Second, to satisfy the assumption of acyclical evolution of phylogenetic models, we excluded identified intra-subtype recombinants from the dataset. While this ensures robust tree inference, it inherently omits the evolutionary contributions of recombinant lineages, which may play a role in the ongoing diversification of the subtype. However, given the adequate temporal signal (correlation coefficient ≈ 0.58) observed across the final datasets, these exclusions are unlikely to alter the broad phylogeographic conclusions. Third, while the reconstruction of transmission routes was robust across methods, the timing of individual introduction events was sensitive to the choice of temporal dating method, with TempEst estimating substantially earlier introduction dates than other methods, particularly for pre-1980 divergence events. Studies drawing conclusions about the precise timing of viral dispersal should therefore consider the uncertainty introduced by temporal method choice.

In conclusion, the global spread of HIV-1 Subtype C is characterized by a dominant, recurring transmission corridor from Africa to Europe, alongside the establishment of independent secondary reservoirs in South America and Asia. This complex dissemination network is primarily sustained by heterosexual transmission, a dynamic frequently obscured by incomplete surveillance annotation but unmasked here through rigorous ancestral trait reconstruction. Methodologically, we demonstrate that while temporal estimates and the inferred timing of individual introduction events vary across dating methods, the reconstruction of global transmission routes and source-sink classifications remains robust across phylogenetic and temporal method combinations. This multi-method benchmarking approach, coupled with concordance-based validation of network centrality and source-sink assignments, offers a scalable alternative to computationally intensive Bayesian frameworks^63^. By providing insight into phylogenetic uncertainty within a manageable timeframe, this strategy serves as a practical, rapid-response pipeline for the real-time genomic investigation of emerging epidemic and pandemic threats.

## Acknowledgements

The opinions expressed in this article are those of the authors and do not reflect the view of the National Institutes of Health, the Department of Health and Human Services, or the United States government.

## Author contributions

X.L. conceived and designed the study. X.L., G.D., A.P., and N.S.T analyzed the data. X.L., S.T., G.D., A.P., and N.S.T interpreted the data and drafted the initial manuscript. All authors reviewed and approved the final manuscript.

## Data Availability

No new data were generated in support of this research. The scripts underlying this article will be shared on reasonable request to the corresponding author.

## Figure Legends

**Supplementary Figure S1. Sampling distributions of five sub-sampled HIV-1 subtype C datasets used in this study.** (A) Stratified by country. (B) Stratified by risk group.

**Supplementary Figure S2. Geographic distribution of five sub-sampled HIV-1 subtype C datasets used in this study.** Points are coloured by risk group, and point size is proportional to the number of genomes sampled from each location. The figure was generated and adapted from Microreact.

**Supplementary Figure S3. Likelihood-mapping analyses of fifteen sub-sampled HIV-1 subtype C datasets.** The corners represent tree-like phylogenetic signals, the sides represent network-like signals, and the central area represents star-like signals corresponding to unresolved phylogenetic structure.

**Supplementary Figures S4–S17. Tangled chain (series of tanglegrams) of phylogenetic trees inferred by four phylogenetic softwares ordered by average-linkage clustering based on ClusteringInfoDistance for fourteen of the fifteen sub-sampled HIV-1 subtype C datasets, excluding locrisk260.1 (Figure 4).**

**Supplementary Figure S18. Statistical benchmarking of 16 phylogenetic-temporal method combinations.** (A) Mean TMRCA estimates varied significantly across temporal but not phylogenetic methods. (B) Mean Evolutionary Rate (ER) estimates were higher for FastTree compared to higher-precision methods. (C) Mean inferred inter-continental introduction counts remained stable across all method combinations. Error bars represent 95% confidence intervals.

**Supplementary Figure S19. Heatmaps of mean TMRCA, mean evolutionary rate, and introduction count variability across 16 phylogenetic-temporal method combinations.** (A) Mean TMRCA: TreeTime produced the earliest dates (∼1927–1931) and treedater the latest (∼1942–1947). (B) Mean evolutionary rate (ER): FastTree produced higher rates than rapid heuristic methods, while TreeTime produced lower rates than other temporal methods. (C) Coefficient of variation (CV) for introduction counts across subsampling replicates, indicating high consistency across all method combinations (CV range: 11.8–16.4%). For all panels, color scale ranges from blue (lower values) to red (higher values).

**Supplementary Figure S20. Temporal distribution of inferred introduction events by dating method.** Kernel density estimates of introduction times across all shared dispersal routes (n = 24,781 events, 58 country pairs), stratified by temporal dating method and aggregated across phylogenetic methods. LSD, TreeTime, and treedater reconstruct two distinct waves of introduction separated by a valley during ∼1975–1985 (gray vertical panel), whereas TempEst infers an earlier first wave (peaking ∼1945) and more continuous low-level dispersal during the intervening period. All methods converge for the second wave (post-1990). Introduction events were filtered to country pairs detected by all 16 method combinations within each dataset.

**Supplementary Table S1. Predicted difficulty of phylogenetic analysis based on multiple sequence alignments using Pythia for each of the fifteen datasets.**

**Supplementary Table S2. Concordance of StrainHub degree centrality rankings across phylogenetic methods and subsampled datasets.** Kendall’s coefficient of concordance (W) was computed for each centrality metric at five resolution levels (continent, country, risk4, risk5, risk6). Three rater groupings were evaluated: phylogenetic reconstruction method (k = 4; FastTree, IQ-TREE, PhyML, RAxML-NG, averaged across datasets and replicates), dataset (k = 5; locrisk260–locrisk574, averaged across phylogenetic methods and replicates), and all method-dataset combinations (k = 60; each unique phylogenetic method × dataset × replicate treated independently). W ranges from 0 (no agreement) to 1 (perfect agreement); significance was assessed using the Friedman chi-squared approximation.

**Supplementary Table S3. Concordance of TreeTime source-sink scores across phylogenetic and temporal dating methods and subsampled datasets.** Kendall’s coefficient of concordance (W) was computed for Export, Import, and Source-Sink Score (SSS) at six resolution levels (continent, region, country, risk4, risk5, risk6). Four rater groupings were evaluated: phylogenetic reconstruction method (k = 4; FastTree, IQ-TREE, PhyML, RAxML-NG, averaged across temporal methods, datasets, and replicates), temporal dating method (k = 4; LSD, TempEst, TreeTime, treedater, averaged across phylogenetic methods, datasets, and replicates), dataset (k = 5; locrisk260–locrisk574, averaged across phylogenetic and temporal methods and replicates), and all method-dataset combinations (k = 240; each unique temporal method × phylogenetic method × dataset × replicate treated independently). At the country level, locations detected in fewer than all 16 method combinations within a given dataset were excluded. W ranges from 0 (no agreement) to 1 (perfect agreement); significance was assessed using the Friedman chi-squared approximation.

**Supplementary Table S4. Linear mixed-effects model results for introduction time benchmarking.** Fixed effect coefficients from a linear mixed-effects model testing the influence of temporal dating method and phylogenetic reconstruction method on estimated introduction times (decimal years). LSD and FastTree serve as reference categories for temporal and phylogenetic methods, respectively. Random intercepts were included for origin country, destination country, transmission route (country pair), and subsampled dataset. ICC = intraclass correlation coefficient; σ² = residual variance; *τ*_₀₀_ = random intercept variance.

