## Supplementary figures and images for "Phylodynamic Analysis of the Global Dispersal and Spatiotemporal Dynamics of HIV-1 Subtype C"

### Fig S1

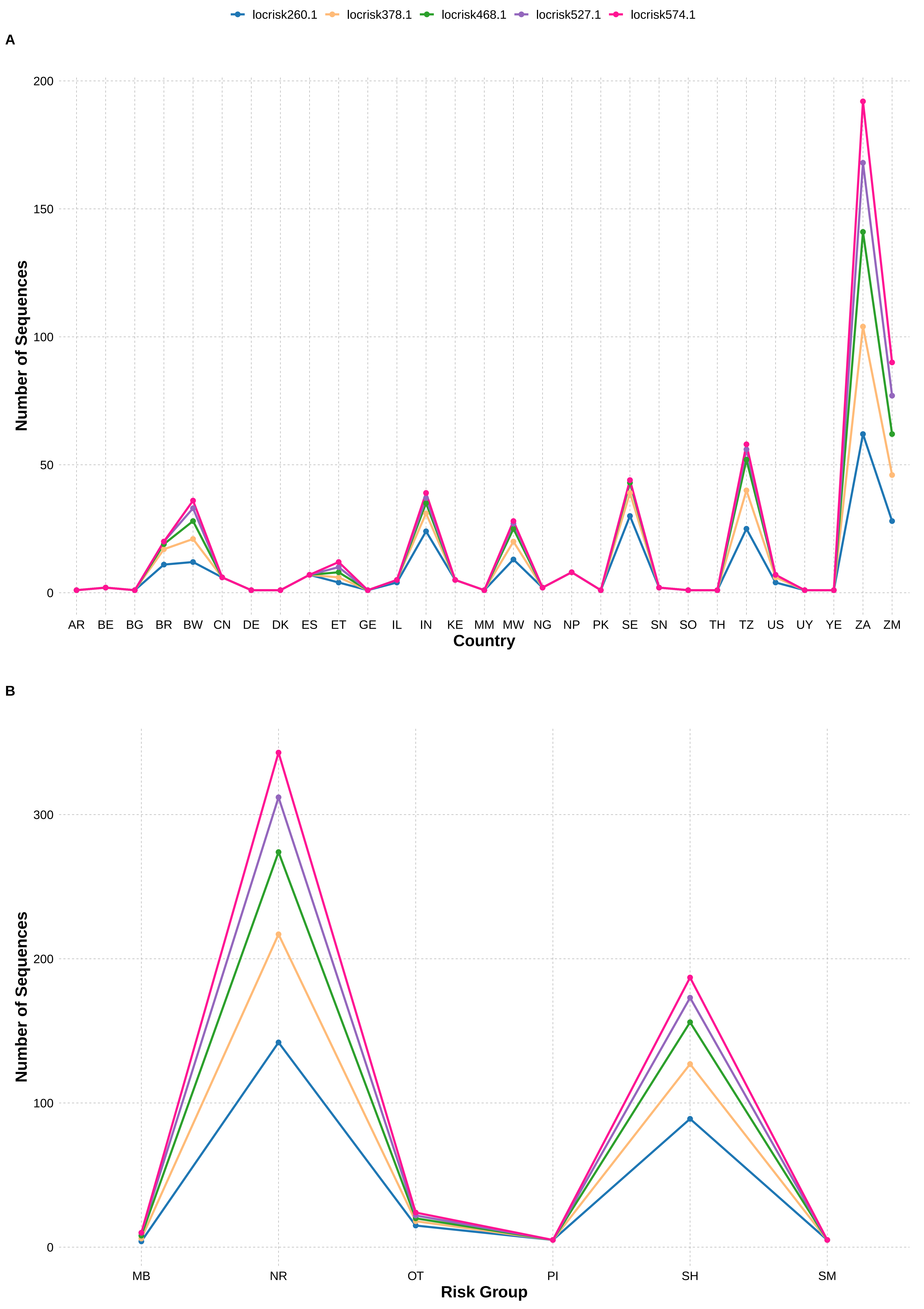

### Fig S2

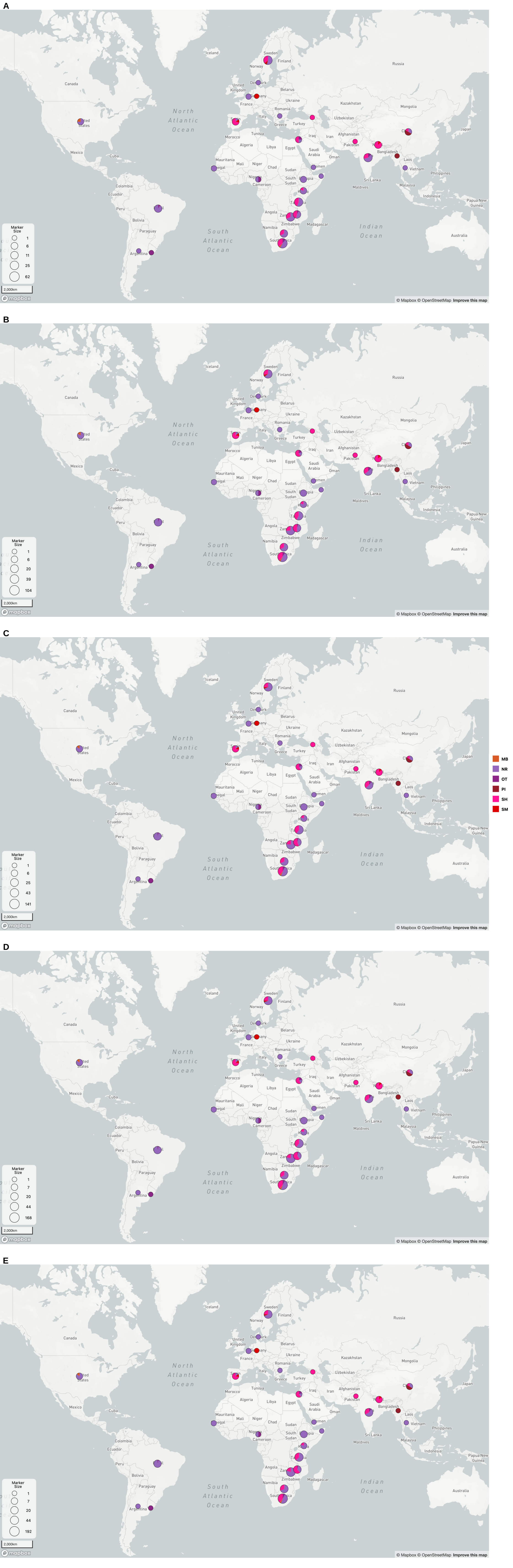

### Fig S3

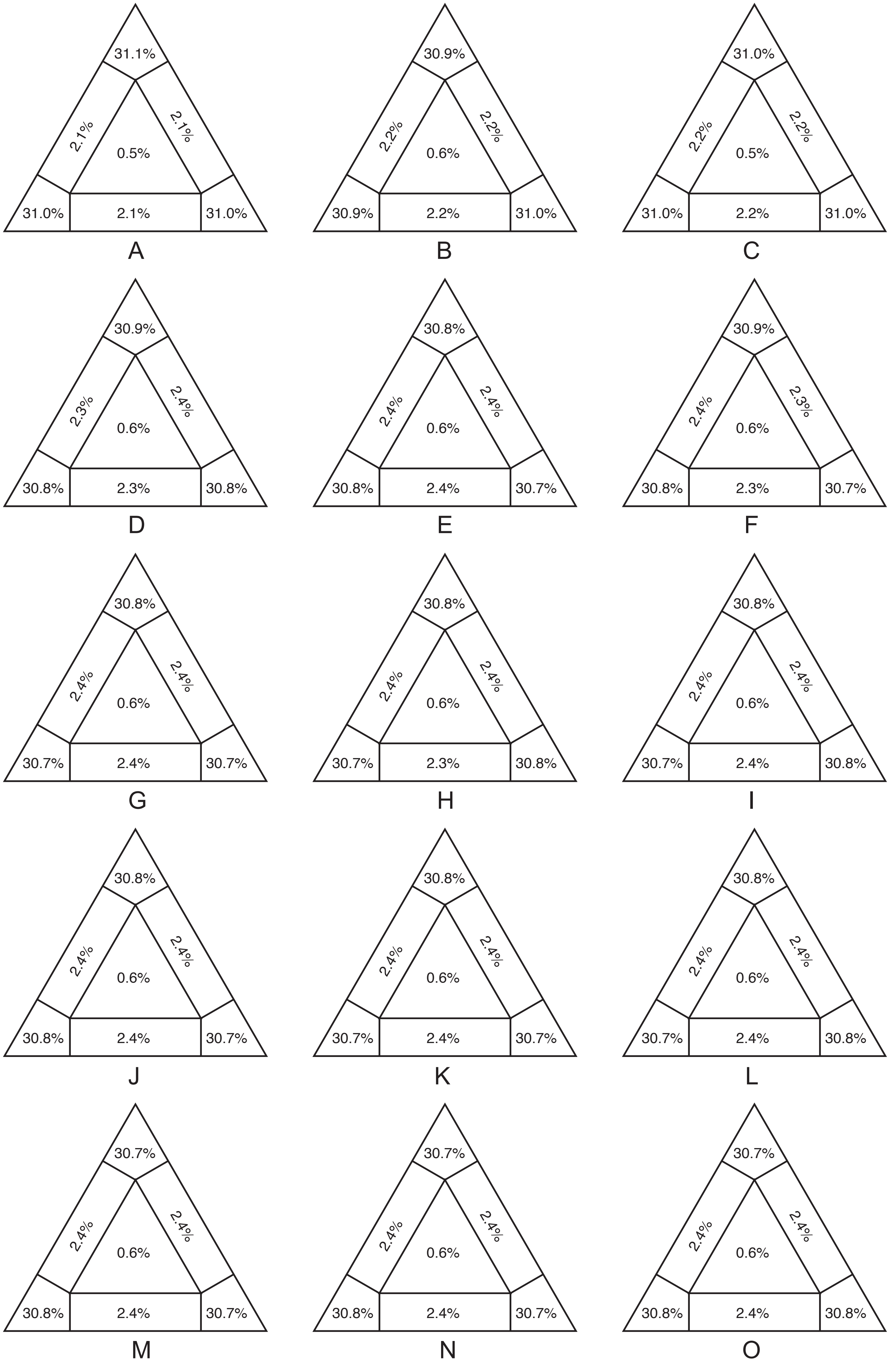

### Fig S4

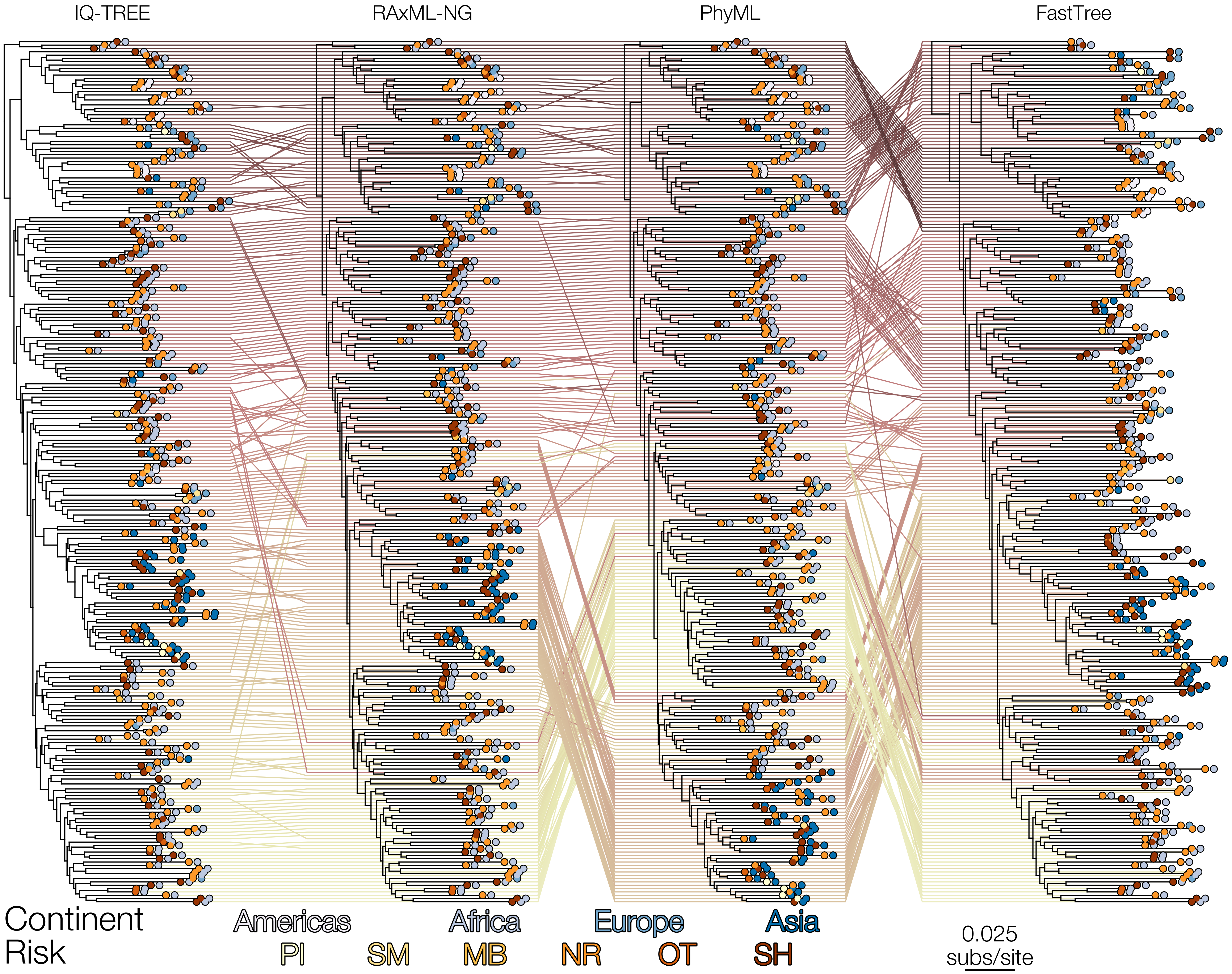

### Fig S5

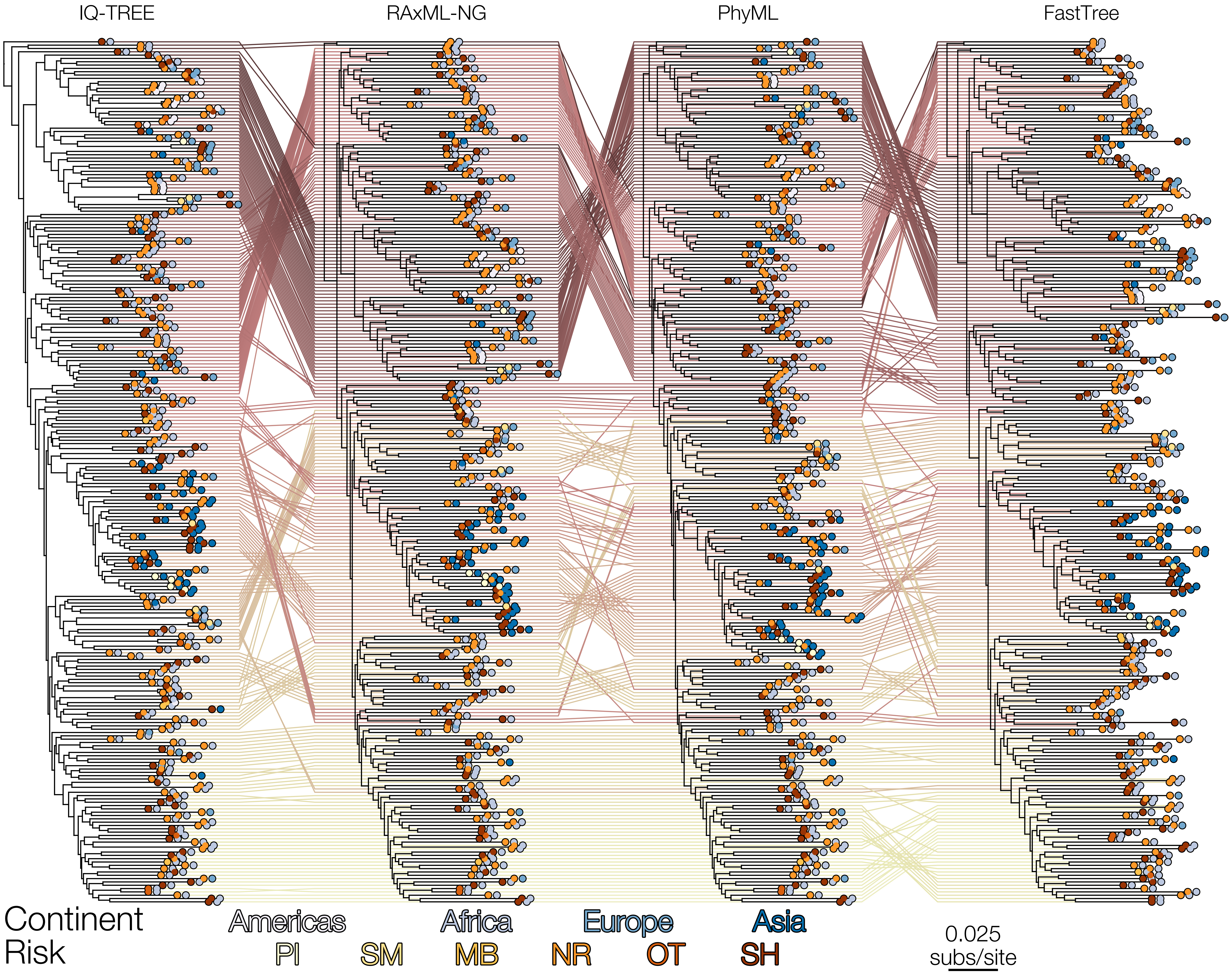

### Fig S6

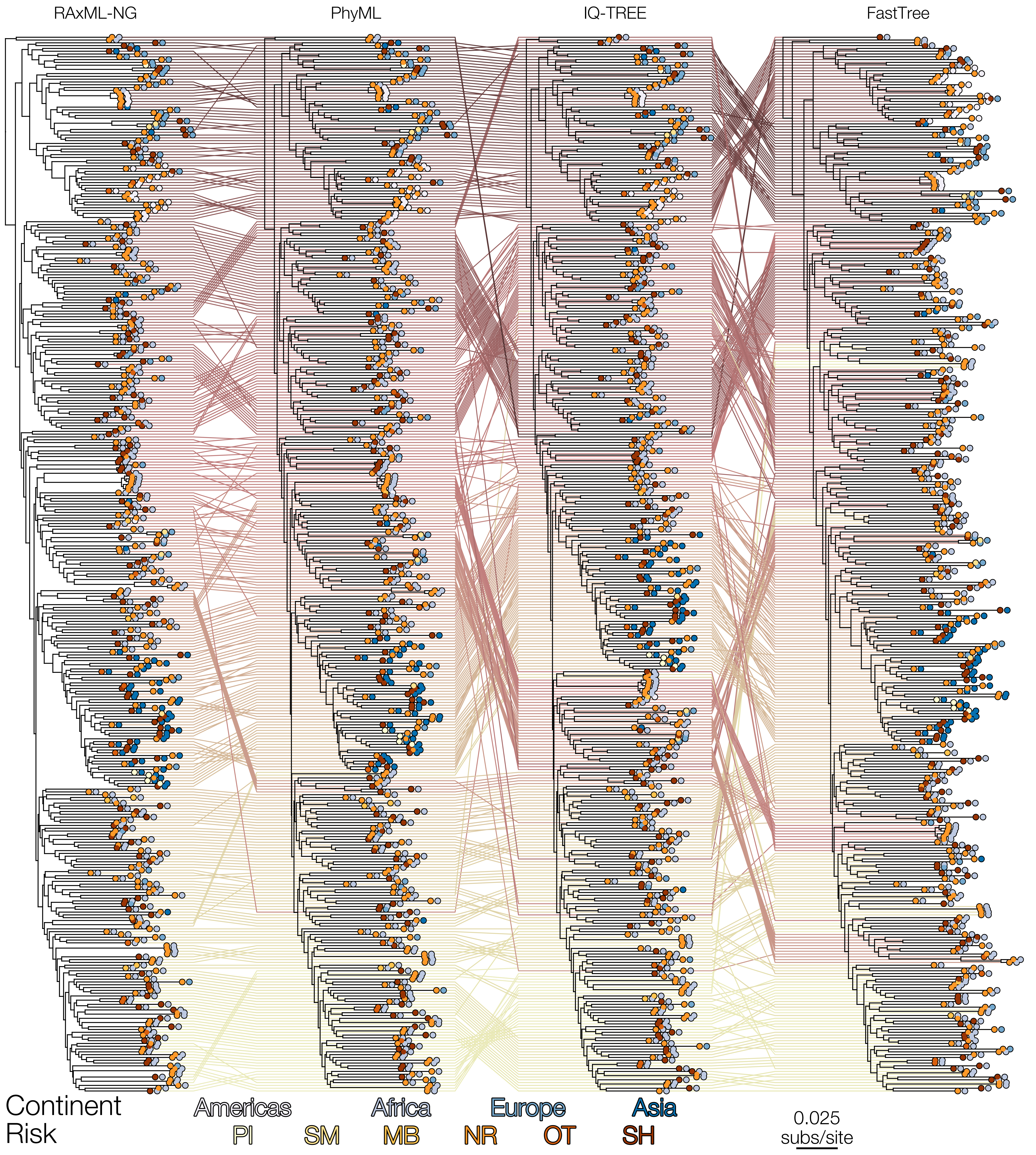

### Fig S7

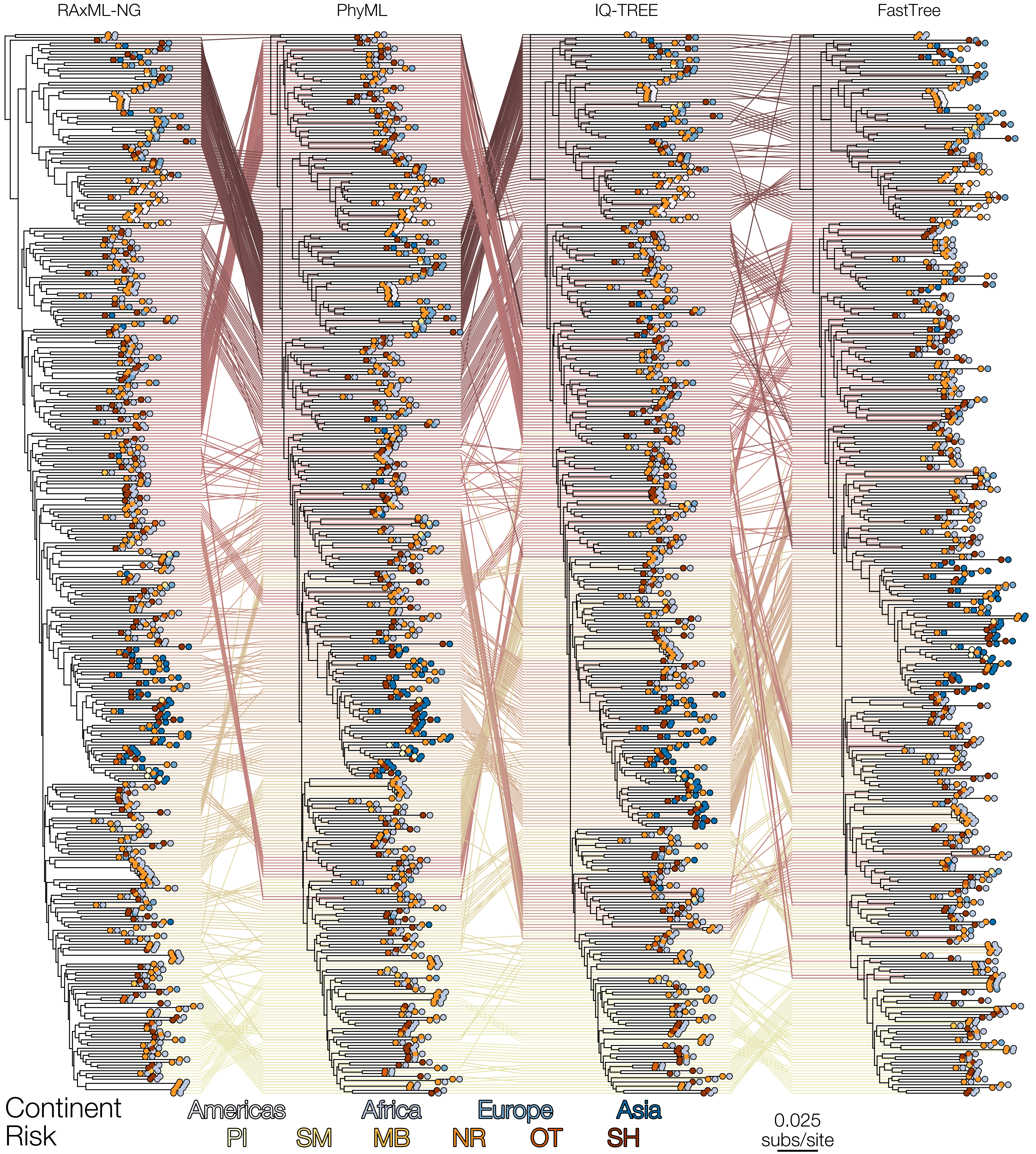

### Fig S8

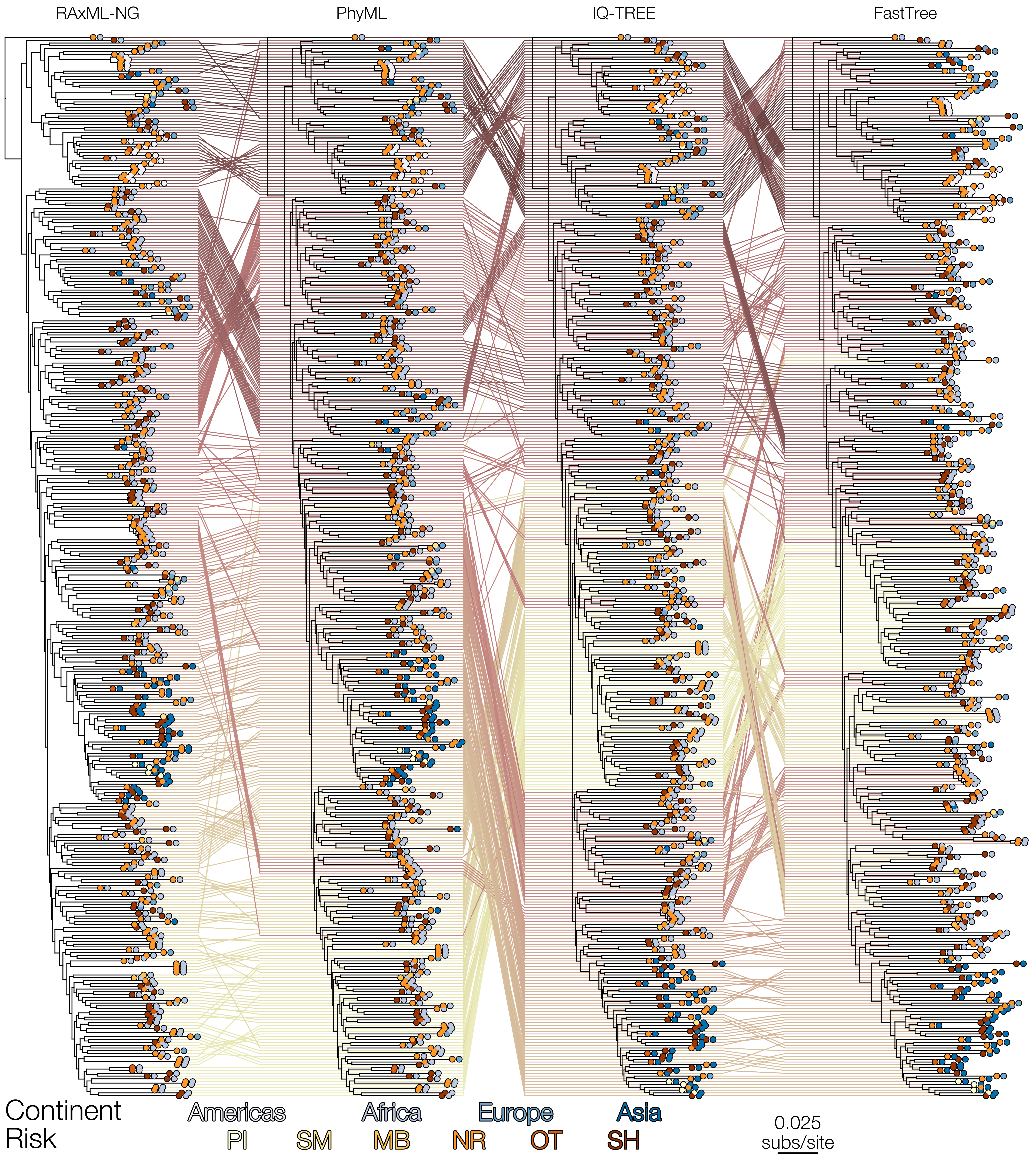

### Fig S9

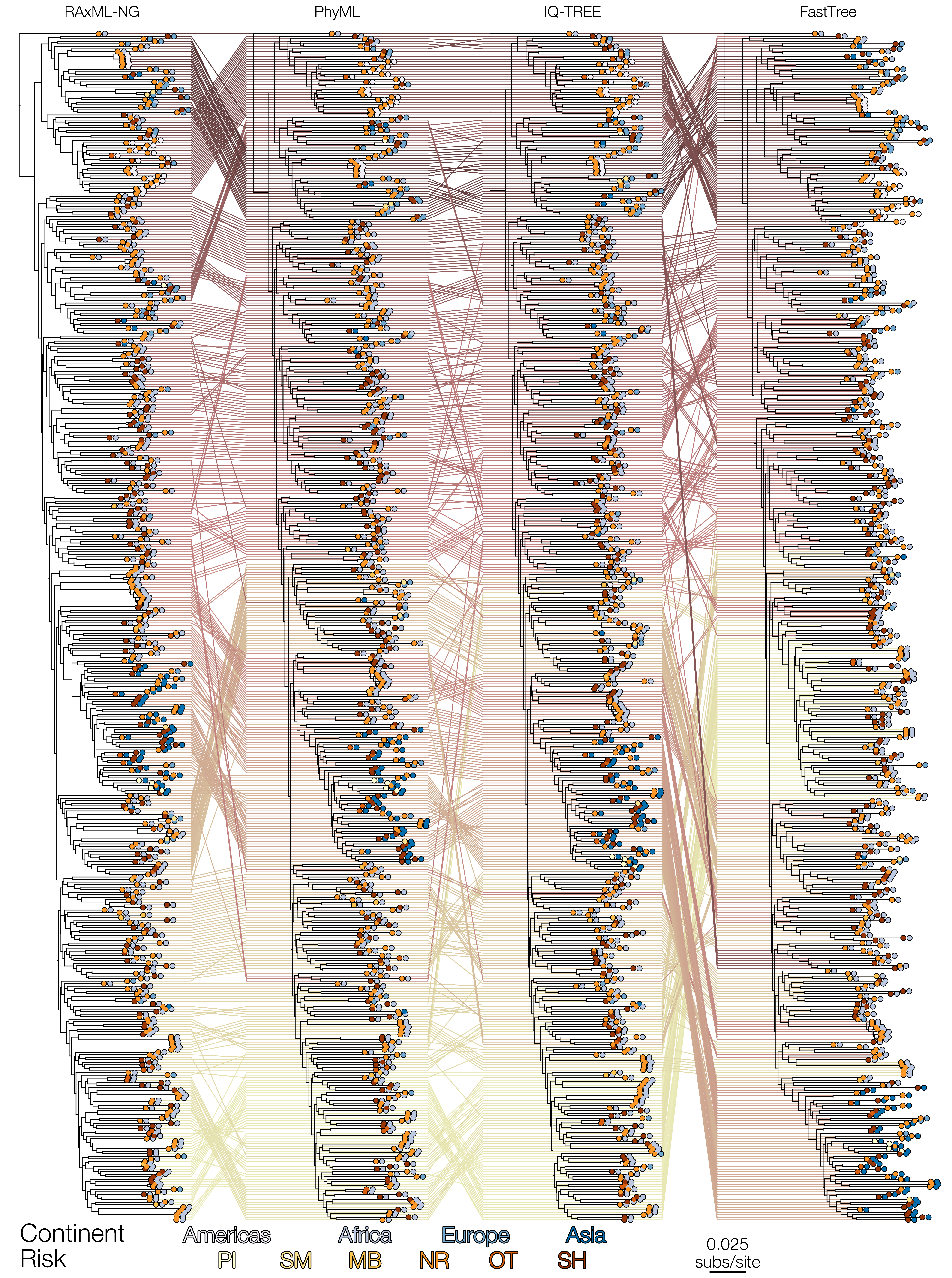

### Fig S10

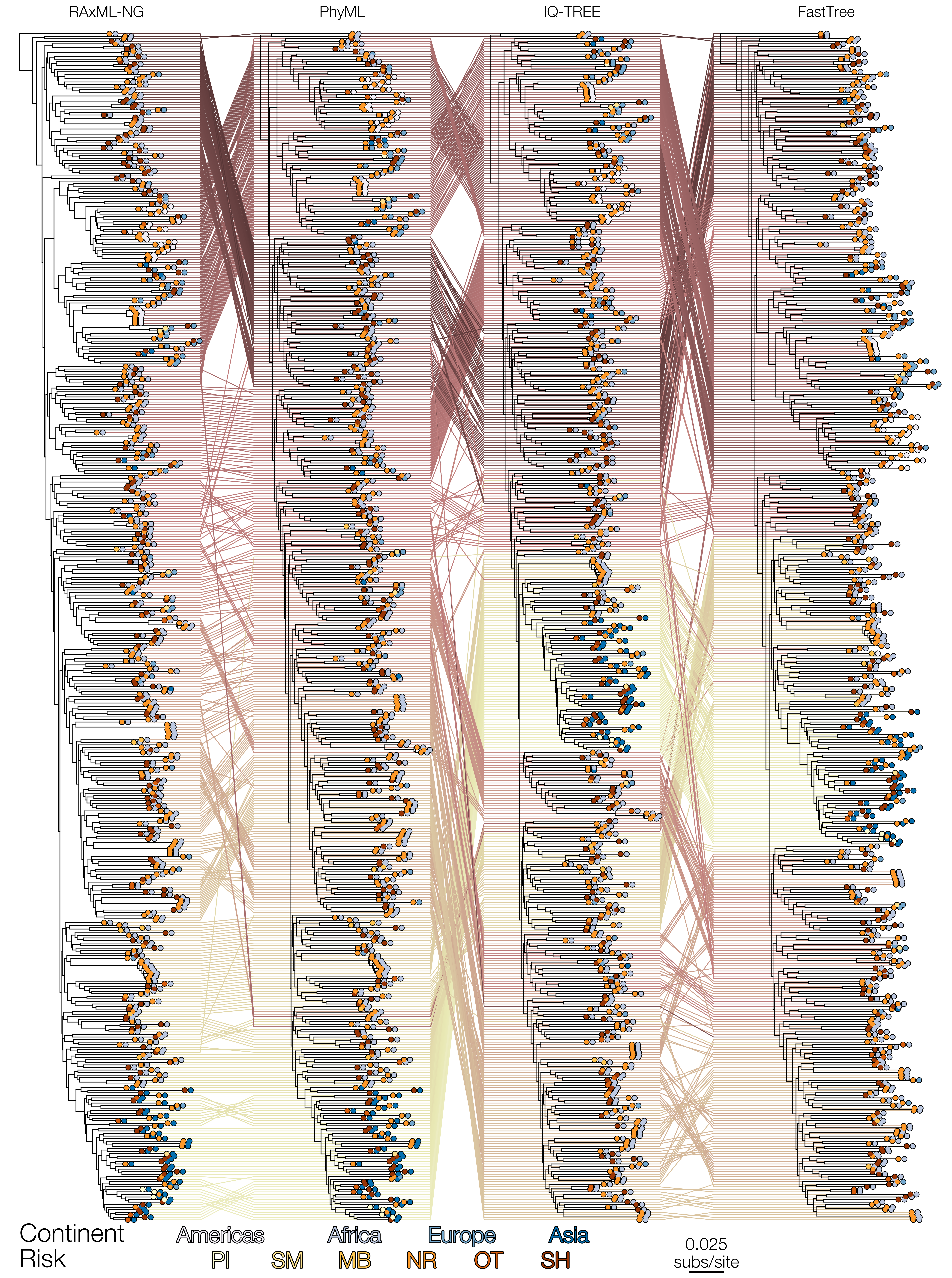

### Fig S11

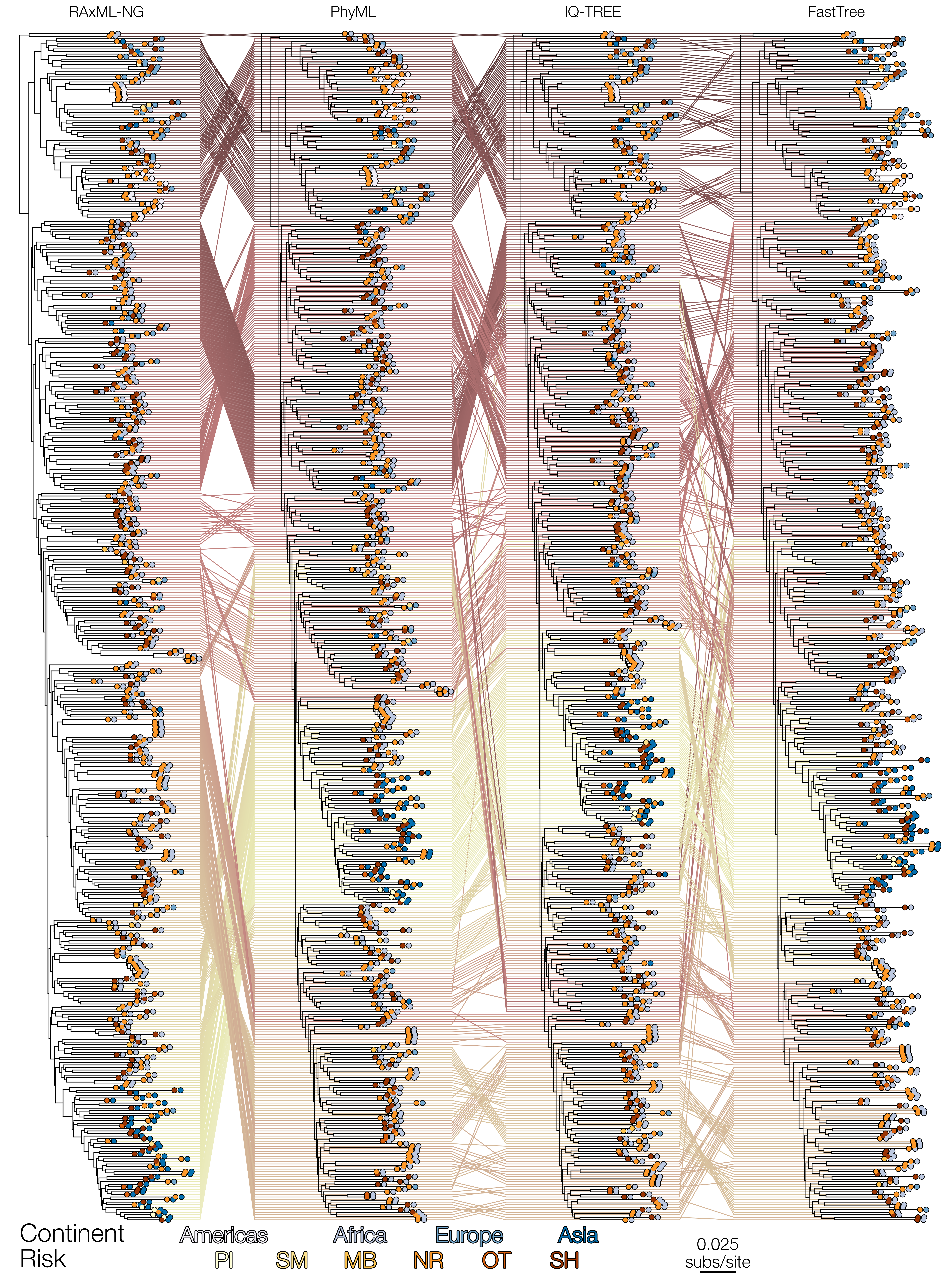

### Fig S12

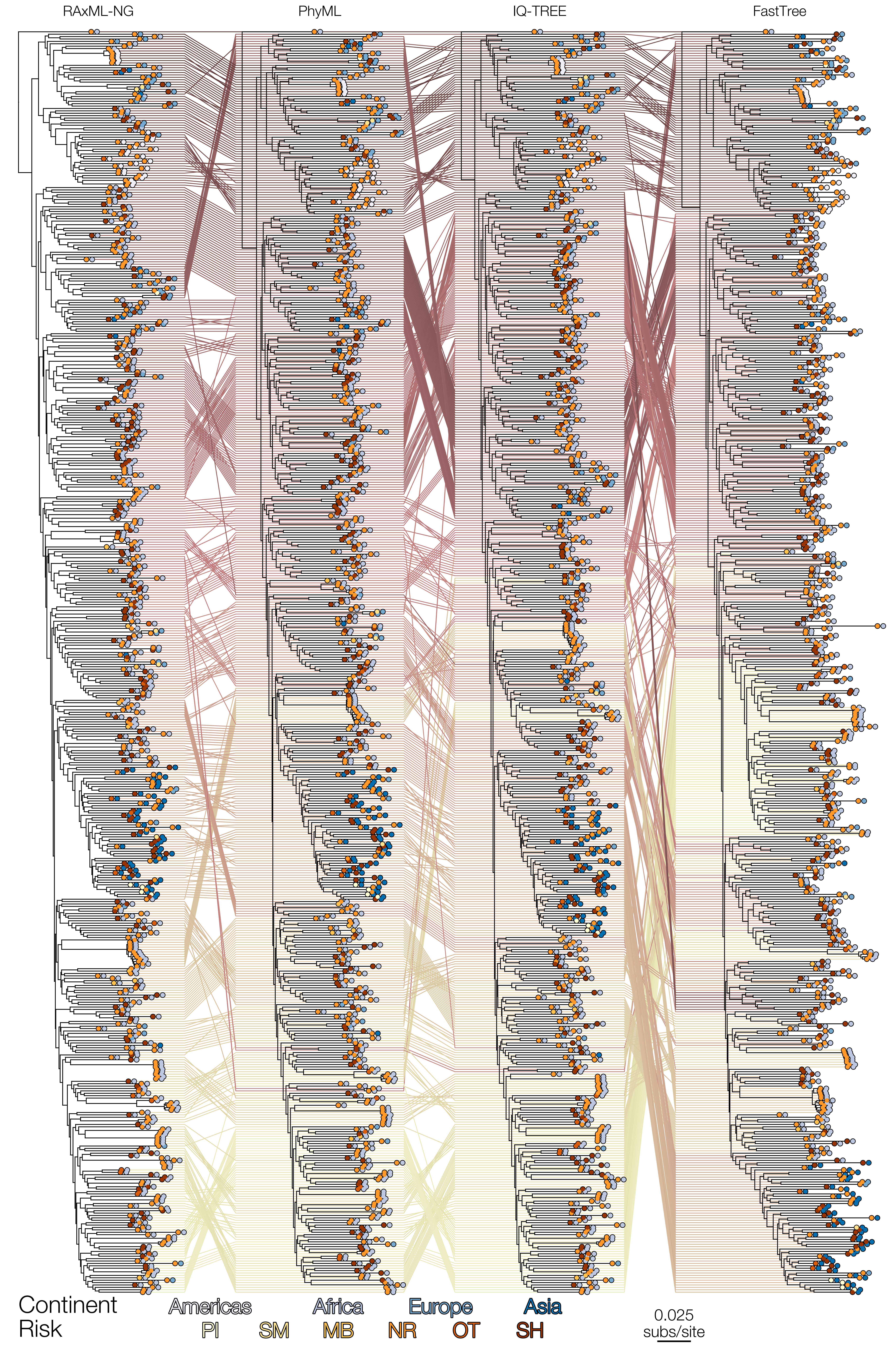

### Fig S13

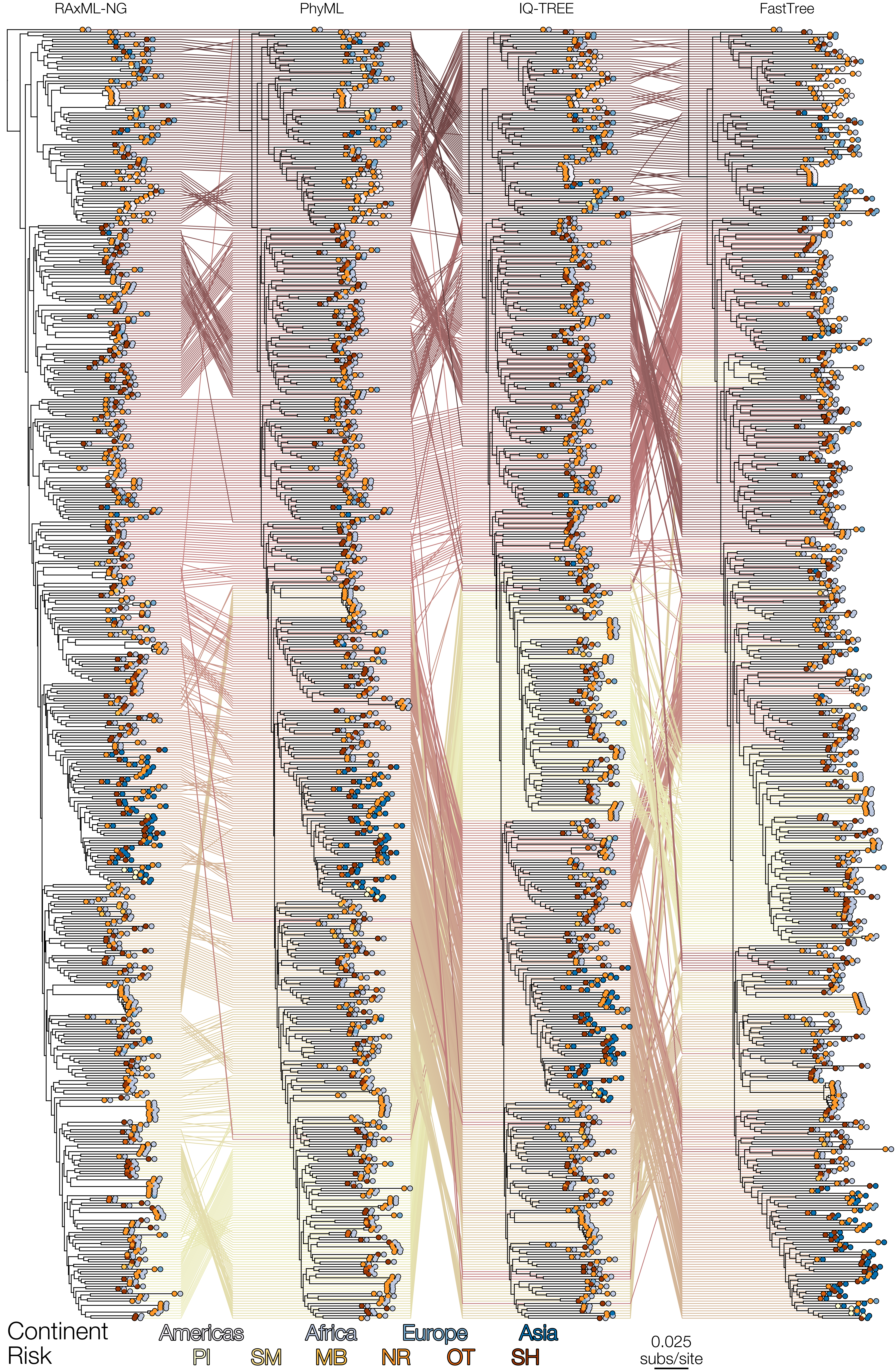

### Fig S14

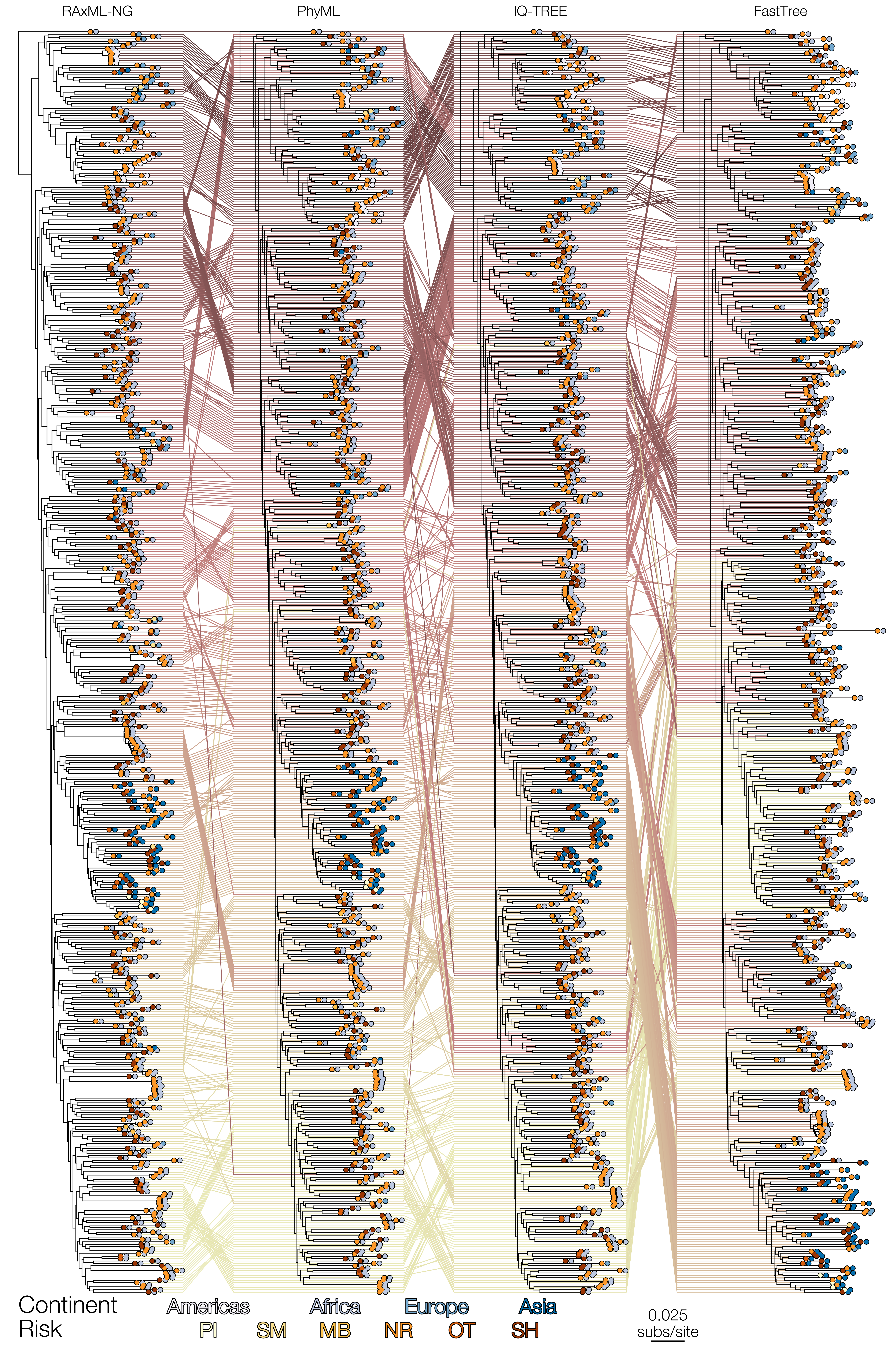

### Fig S15

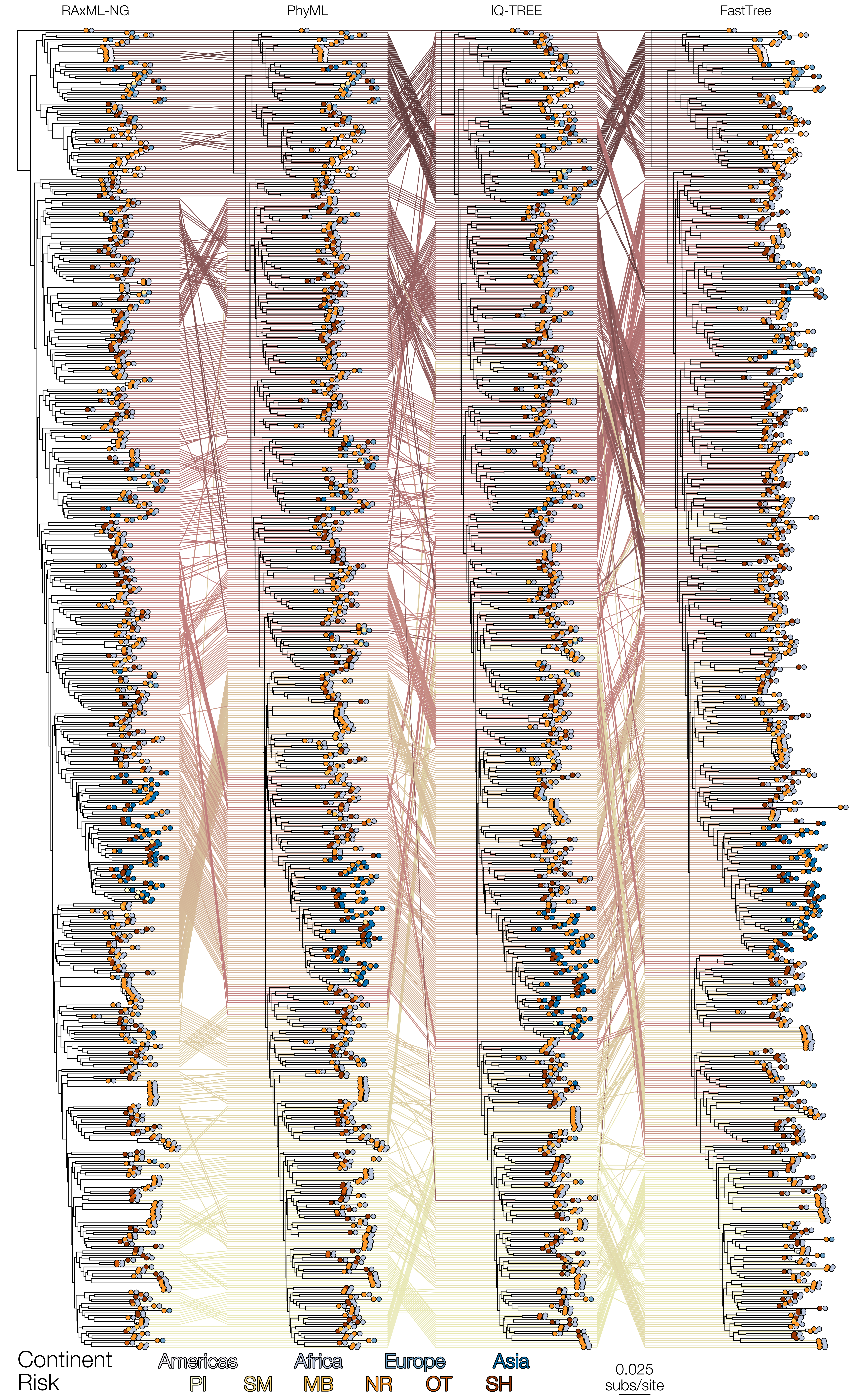

### Fig S16

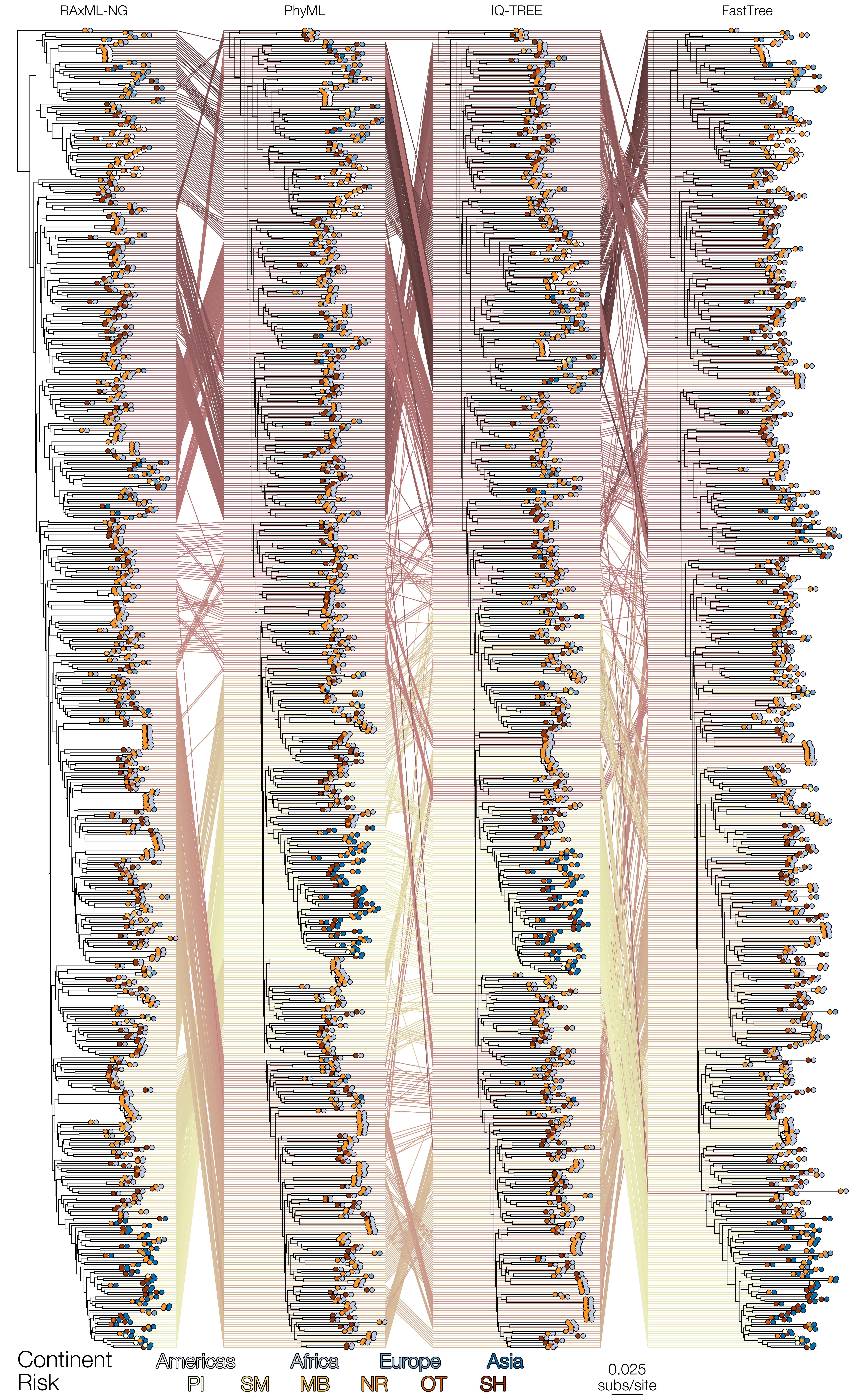

### Fig S17

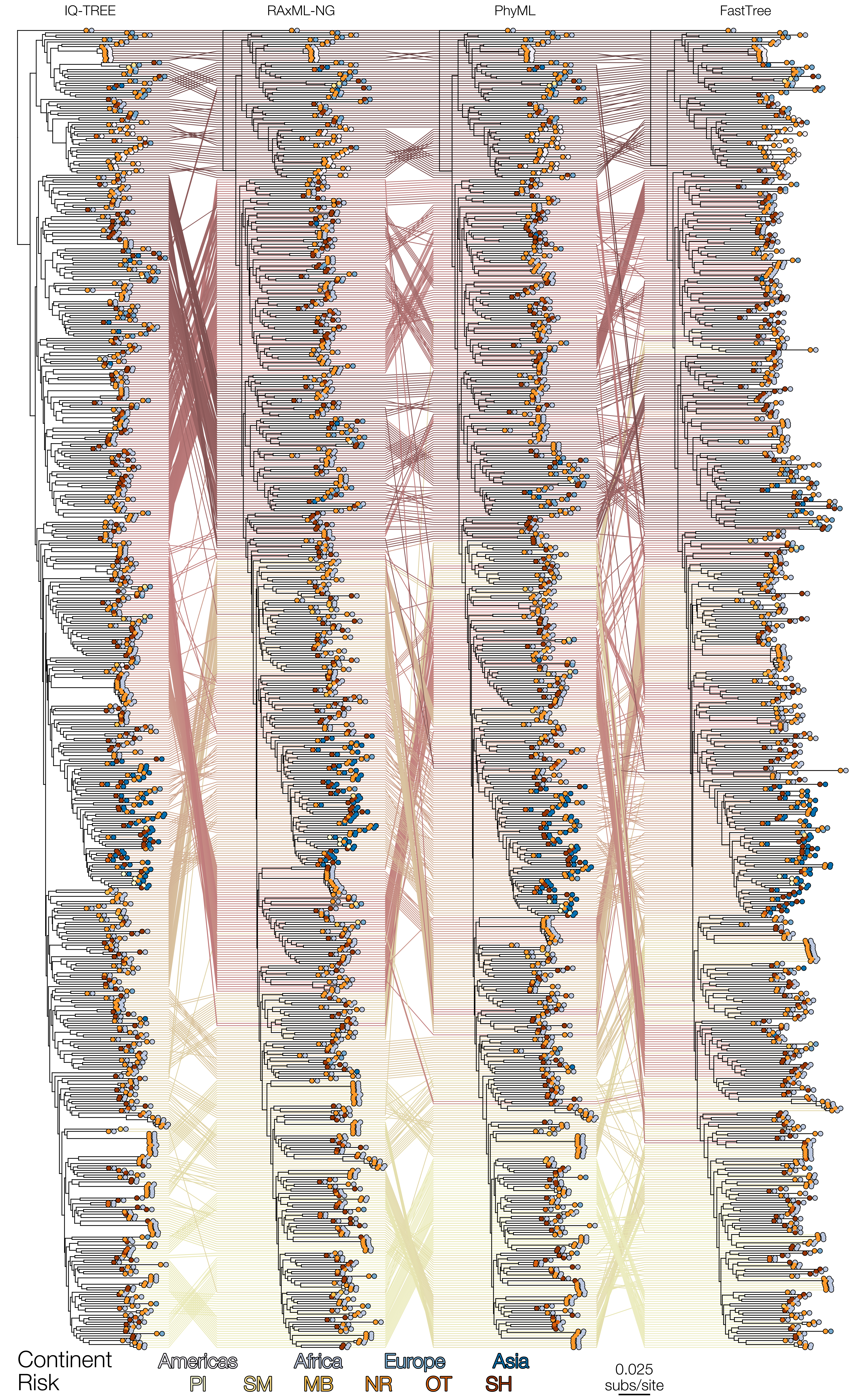

### Fig S18

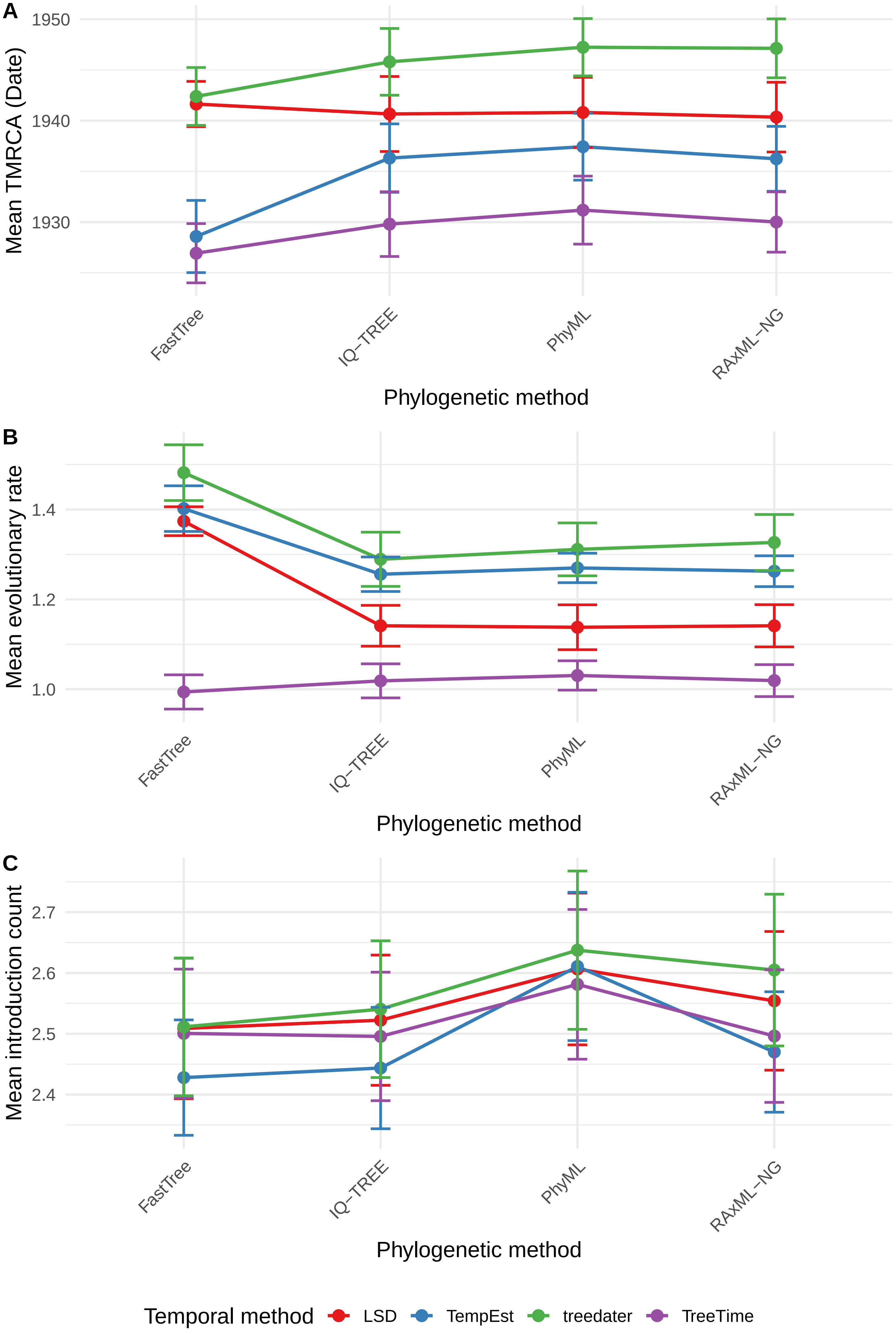

### Fig S19

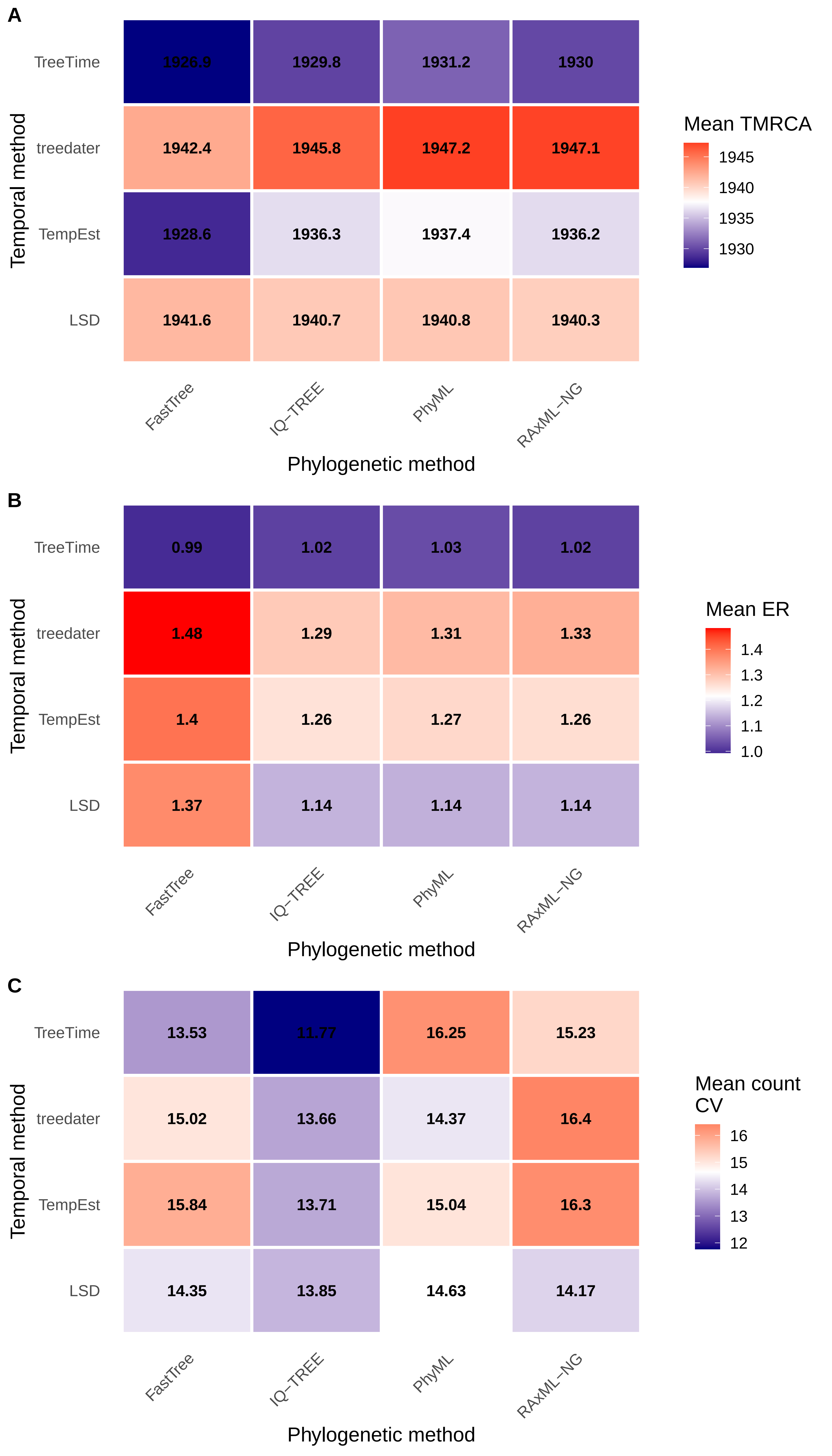

### Fig S20

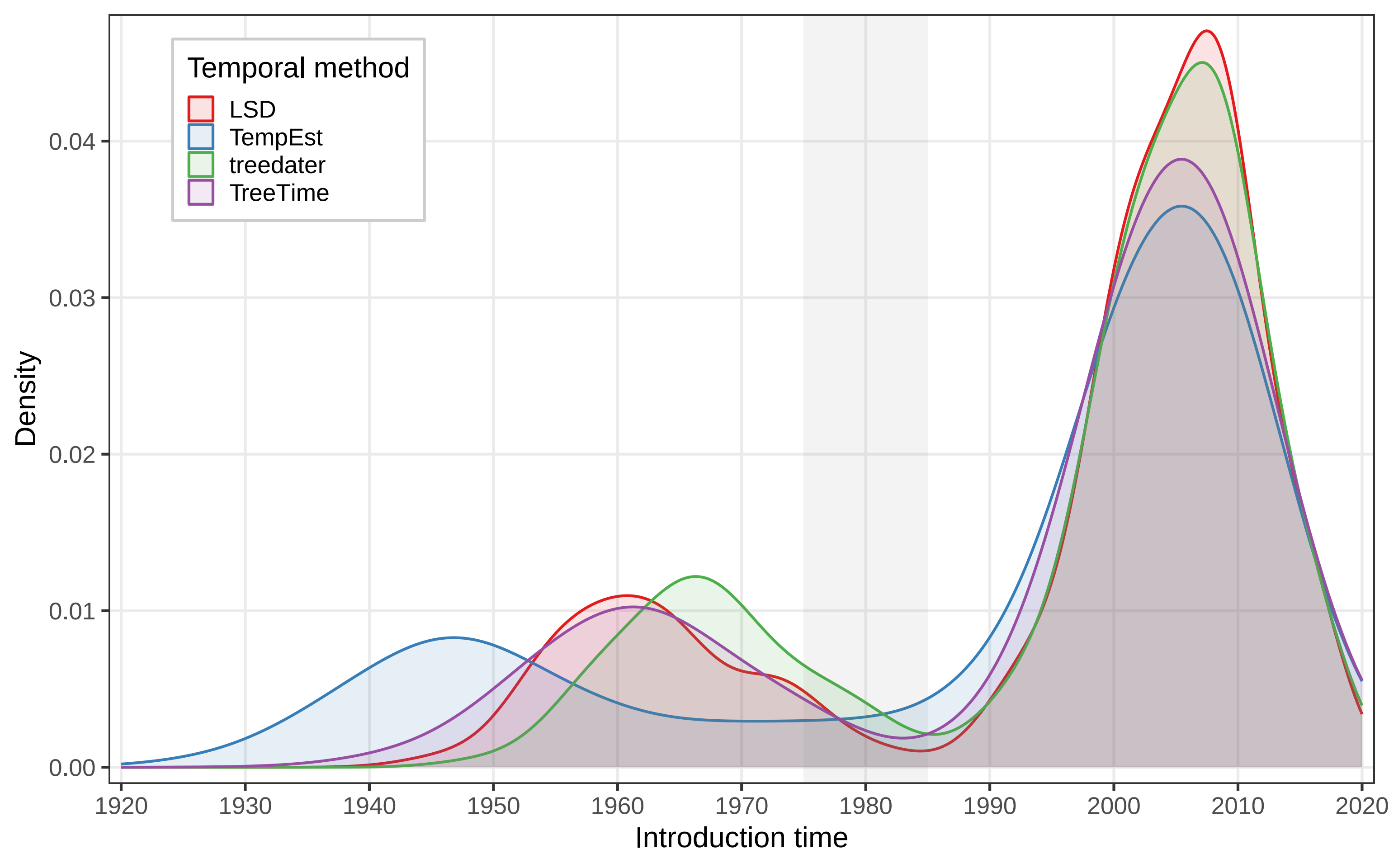
